# Phosphoglucomutase A mediated regulation of carbon flux is essential for antibiotic and disease persistence in *Mycobacterium tuberculosis*

**DOI:** 10.1101/2024.06.27.600960

**Authors:** Taruna Sharma, Shaifali Tyagi, Rahul Pal, Jayendrajyoti Kundu, Sonu Kumar Gupta, Vishawjeet Barik, Vaibhav Kumar Nain, Manitosh Pandey, Prabhanjan Dwivedi, Bhishma Narayan Panda, Yashwant Kumar, Ranjan Kumar Nanda, Samrat Chatterjee, Amit Kumar Pandey

## Abstract

The long-term survival of Mtb mandates judicious utilization of the available resources inside the host. Uninterrupted access to host-derived nutrients holds the key to the success of Mtb. Phosphoglucomutase enzyme besides synthesizing glycogen, which serves as a nutrient reservoir, also helps modulate the carbon flux in different pathogens. Studies on the role of glycogen metabolism in disease progression, reactivation, and drug susceptibility in tuberculosis are severely lacking. To investigate this, we generated an Mtb strain (Δ*pgmA*) devoid of the gene that encodes for the enzyme phosphoglucomutase A (*pgmA*). The absence of *pgmA* impedes the ability of the pathogen to survive under nutrient-limiting and reactivation conditions. In the current study, we demonstrate that the absence of cell membrane-associated glycolipids in Δ*pgmA* compromised the cell wall integrity and increased the susceptibility of Δ*pgmA* to various stresses. Interestingly, in comparison to the wild type, low cAMP levels in Δ*pgmA* imparted an enhanced growth phenotype on cholesterol. Differential gene expression and carbon flux analysis suggest that stored carbon in the form of glycogen is essential for the survival of Mtb under nutrient-limiting conditions. Finally, we demonstrate that the *pgmA* gene of Mtb is essential for the growth of Mtb inside the host. Overall, this study unveils the significance of *pgmA-*mediated regulation of membrane glycolipids and its implication on antibiotic and disease persistence in tuberculosis. Additionally, information derived from this study will help design anti-TB strategies that are novel, short, and more efficient.

## Introduction

Metabolic plasticity is crucial for pathogens to adapt to changing environments and uphold structural stability despite varying nutrient availability. Consequently, in Mtb, an obligate intracellular pathogen that is completely dependent on the host, metabolic plasticity help plays a dominant role in the survival of the pathogen inside nutrient-deprived intracellular niches inside the host ^1^. To counter this, Mtb has evolved strategies that regulate the carbon flux for an efficient utilization of available resources. The continuous metabolic shift between growth and biosynthesis is tightly regulated by a network of metabolic and signalling pathways that facilitate this process. Despite playing a major role in the disease pathogenesis, the precise mechanism that maintains this balance remains largely elusive. Phosphoglucomutase (*pgmA*), an enzyme that reversibly converts Glucose-6-phosphate (G6P) to Glucose-1-phosphate (G1P), is one such enzyme that regulates the flux of the carbon by modulating this transition ^2, 3^. As and when the need arises, this reversible transformation enables the pathogen to regulate the flow of carbon between various catabolic and biosynthesis pathways. The catabolic pathway originating from G6P is known to support growth by providing energy. Alternatively, during biosynthesis, the downstream metabolites generated using G1P essentially contribute towards the synthesis and maintenance of the lipid-rich cell wall. However, *pgmA*-dependent regulation of the central carbon metabolism (CCM) and its role in tuberculosis disease biology remains relatively unexplored.

In certain bacterial species, the inhibition of the *pgmA* gene has been associated with decreased energy production and diminished bacterial survival, particularly during shifts in carbon flux within the host environment ^4, 5^, cell wall biosynthesis, and antibiotic resistance ^6, 7, 8, 9, 10, 11^. Non-pathogenic Δ*pgmA* strains from *Brucella melitensis* and *Streptococcus iniae* have shown *pgmA* to be a promising candidate for the development of attenuated vaccine strains ^12, 13^. Metabolic products from *pgmA* dependent G1P arm includes crucial intermediates such as trehalose, UDP-glucose, maltose-1-phosphate and finally glycogen whose role in membrane biogenesis and integrity is well known ^14, 15, 16, 17, 18^. Glycogen is a universally conserved storage molecule, is known to act as a carbon reservoir during nutrient scarcity ^19^. In *Mycobacterium smegmatis* (Msm), it was reported that in addition to the role of glycogen as a conventional storage macromolecule, a continuous synthesized and degradation of glycogen throughout the exponential growth phase happens suggesting its involvement as a carbon capacitor for glycolysis during active growth ^20^. Although we have some information on the role of glycogen metabolism in the growth and survival of Msm, its role in mycobacterial drug susceptibility, virulence, and pathogenesis is lacking.

In this study, we aimed to elucidate the role of *pgmA* in maintaining carbon flux balance during the nutrient scarcity model in Mtb. To accomplish this, we generated a genetically modified Mtb strain lacking the *pgmA* gene (Δ*pgmA*). We found that relative to the wild type, Δ*pgmA* demonstrated a decrease in their ability to grow under both nutrient-limiting and reactivation conditions. The absence of *pgmA* in Mtb resulted in reduced generation of glycogen, trehalose, and other membrane-associated glycolipids compromising both the cell wall integrity and the ability to survive under stress. Intriguingly, we found that *pgmA*-mediated growth modulation on cholesterol is one of the critical drivers of antibiotic and disease persistence in tuberculosis. Understanding the intricacies of central carbon metabolism in mycobacteria holds potential for innovative therapeutic strategies aimed at disrupting their energy equilibrium and virulence. Furthermore, targeting mycobacterial *pgmA* could provide a promising approach for developing tailored antimicrobial agents to combat mycobacterial infections.

## Results

### *pgmA* is essential for survival of Mtb under nutrient limiting condition

*pgmA* is an isomerase involved in the inter-conversion of G1P to G6P (Fig. 1A), a critical step involved in the synthesis of metabolites fundamental to central carbon metabolism (CCM). To further characterize this gene we generated *pgmA* gene deletion mutant strain in H37Rv by homologous recombination technique (Fig. S1A). Further, the gene deletion in Δ*pgmA* at the gene and transcript level was confirmed by PCR and RT-PCR methods respectively (Fig. S1B and S1C). Additionally, as a control, we also generated the complemented strain by expressing the *pgmA* gene in the mutant strain. Since glycogen is the final product of the pathway, we initially measured its levels. As expected, we observed a significant decrease in the level of glycogen in the Δ*pgmA* relative to the wild type which was restored in the complemented strain (Fig. 1B, also see Fig. S2A). Despite altered glycogen level, we did not observe any difference in the ability of Δ*pgmA* to grow under nutrient-sufficient conditions (Fig.1C). Since glycogen is universally known to act like a nutrient reservoir across, we were interested to know its role in supporting the growth of Mtb under both nutrient limiting and resuscitation conditions (Fig. 1D). Interestingly, relative to the wild type and the complemented strains, the growth of Δ*pgmA* was found to be impaired when exposed to starvation condition observed in terms of both colony forming unit (Fig. 1E) and total biomass (Fig. S2B). Additionally, the reactivation of the starved Δ*pgmA* in enriched media also demonstrated a delayed resuscitation phenotype in comparison to the wild type and the complemented strains (Fig.1F). Lower ATP levels were observed under starvation conditions indicating compromised fitness and low metabolic activity of Δ*pgmA* in such environment (Fig. S2C). The above observation suggests that *pgmA* is crucial for the glycogen biosynthesis pathway and facilitates the growth of Mtb under both nutrient-limiting and resuscitation conditions.

**Figure 1:**
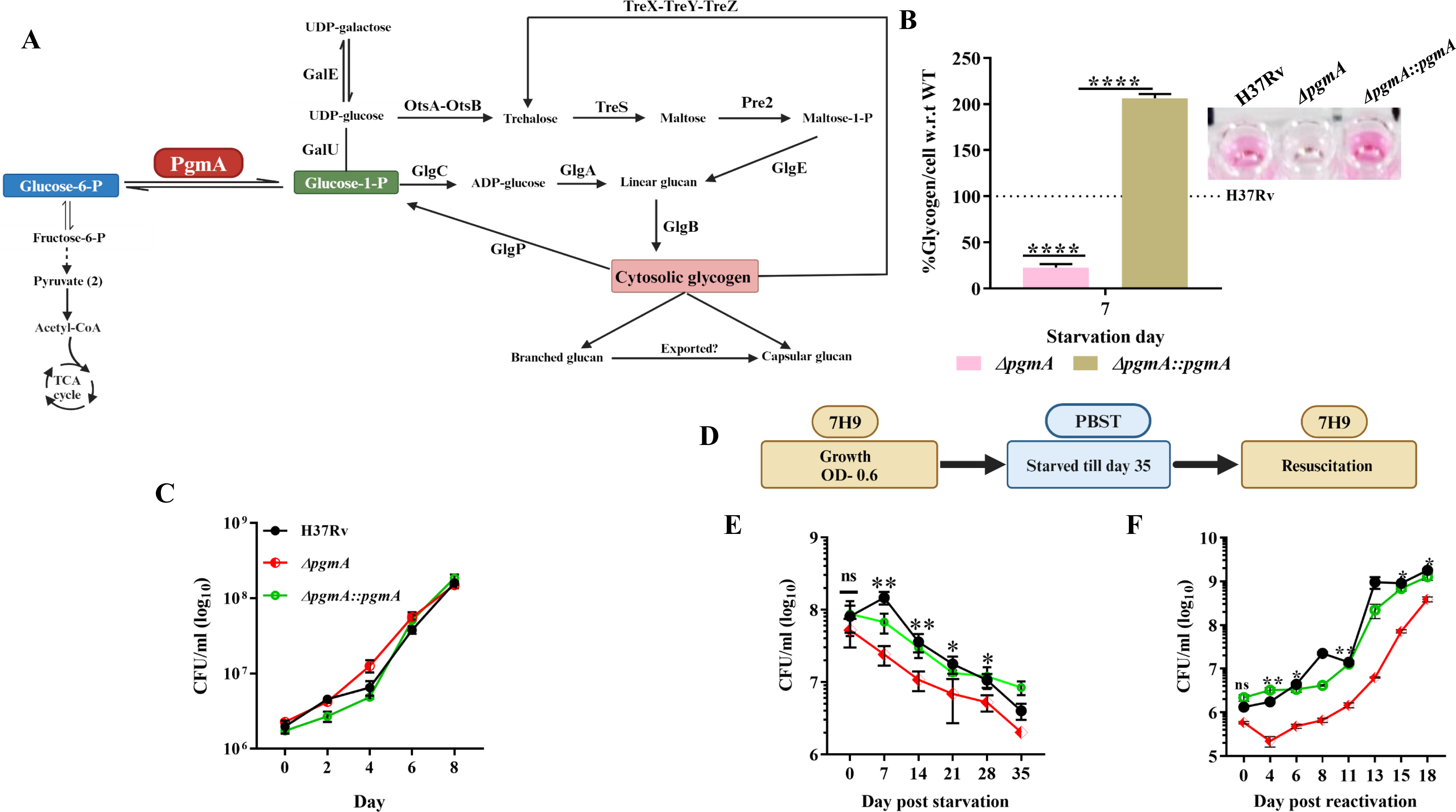
*pgmA* is essential for survival of Mtb under nutrient limiting condition. **(A)** Schematic representation of *pgmA* catalyzing the inter-conversion of G6P and G1P depicting the intermediates crucial for biosynthesis and growth. (B) Altered glycogen levels in the absence of *pgmA* at day 7 starvation in PBST (also see; Fig. S2A). The inset shows the calorimetric estimation of glycogen levels, highlighting visual differences between the strains. (C) Growth kinetics of Δ*pgmA* under standard growth conditions (D) Illustration outlining the experimental approach for carbon starvation and subsequent resuscitation (E) Δ*pgmA* fails to survive under carbon starvation and is essential during (F) resuscitation. Statistical significance (B) to (C) and (E) to (F) was determined using unpaired, non-parametric two-tailed t-test *P≤0.05 and **P ≤ 0.005. Data represent mean ± SEM (standard error mean) for technical triplicates.

### *pgmA* is crucial for maintaining cellular architecture of Mtb

*pgmA-*dependent synthesis of G1P acts as a precursor of the biosynthesis of critical metabolites essential for the synthesis and transportation of cell wall-associated sugars and lipids ^21, 22, 23, 24^. We hypothesize that the absence of *pgmA* will significantly compromise the membrane integrity of the Mtb. The same was first tested by performing the ethidium bromide (EtBr) permeability assay. High EtBr uptake was observed by Δ*pgmA* relative to the wild type and the complemented strain (Fig.2A) suggests that the absence of *pgmA* enhances the cell permeability by altering the integrity of the cell membrane. Further, total lipids extracted from cultures were separated using thin-layer chromatography (TLC) technique. Total lipid was separated on a silica plate (stationary phase) using different combinations of the organic solvents (mobile phase) (Fig. S3A). Surprisingly, TLC analysis of membrane lipids revealed a significant reduction in band intensity of cell wall-associated lipids including triacyl-glycerol (TAG), PDIM (Fig 2B and Fig. S3B), mycolic acids (Fig. 2C) in Δ*pgmA* relative of the wild type.

**Figure 2:**
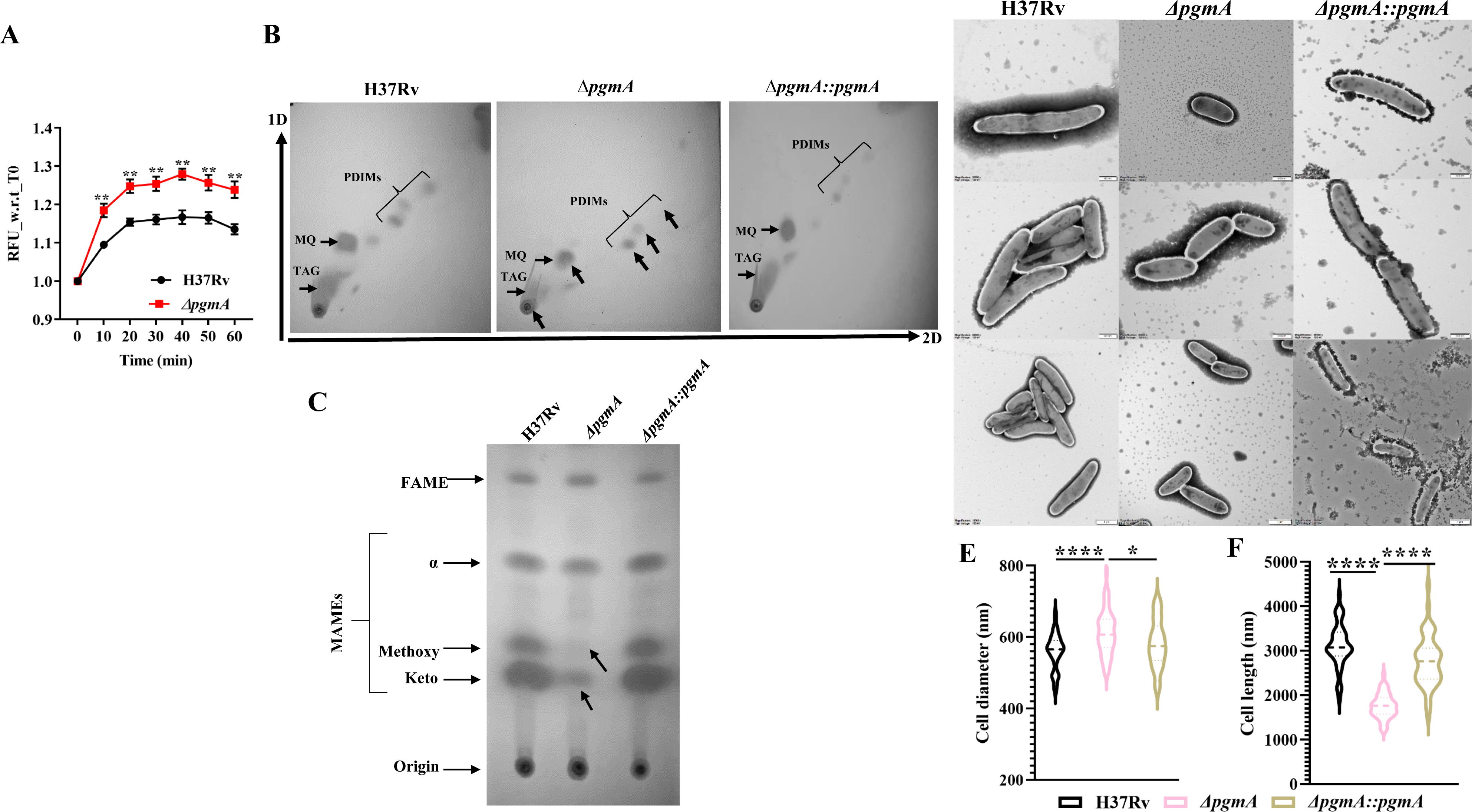
*pgmA* is crucial for maintaining cellular architecture of Mtb. **(A)** Increased membrane permeability observed in the Δ*pgmA* strain. In (B) to (C) total lipids were separated by growing bacterial cells till A_600_ reaches 1-1.5. TLC reveals deficiencies in (B) PDIM-A, TAGs, and (C) mycolic acid in Δ*pgmA*. 2D lipid separation was achieved using hexane:ethyl acetate (98:2) for the first dimension and hexane:acetone (98:2) for the second dimension. In (D), TEM images post seven days of starvation depict (E) an increase in overall cell diameter and (F) a decrease in cell length (*n=50* cells each strain) of Δ*pgmA*. The cells were visualized at 20000X to 12000X magnification (top to bottom) on 500nm to 1µm scale (top to bottom) at 120kV. In (E) and (F), the cell size measurement was done using RADIUS 2.0 EMSIS software. Statistical significance (A), (E) and (F) was determined using unpaired, non-parametric two-tailed t-test *P≤0.05 and ****P ≤ 0.00005. Abbreviations: PDIM = phthiocerol dimycocerosates, TAGs= Triacyl glycerol, FAMEs= Fatty acid methyl ester MAMEs= Mycolic acid methyl ester, α=alpha, MQ= Menaquinone,.

To understand the morphological implications of these lipid abnormalities on Δ*pgmA*, we visualized these strains using TEM (Fig. 2D). We found that the absence of the *pgmA* gene resulted in a shortening of the overall length (Fig.2E) with a simultaneous increase in the diameter of individual bacterium (Fig.2F). This resulted in imparting a more spherical shape to the mutant relative to the elongated morphology observed in the wild type and complemented strain. Interestingly, in comparison to the wild type and complemented strains, defects in overall membrane lipid biosynthesis significantly impacted the ability of Δ*pgmA* to form biofilm (Fig. S2C). The lipid isolated from the biofilms suggests effect was found to be associated with a deficiency in PDIMs and TAGs (Fig S2D). This observation was linked by evaluating the MIC of vancomycin wherein, as expected, the Δ*pgmA* strain exhibited a significantly reduced IC90 in response to the drug treatment (Fig. S3E). The observational data presented above underscores the pivotal role of *pgmA* in preserving the integrity of the membrane and the overall cell structure of Mtb.

### Deletion of *pgmA* renders Mtb susceptible to various stresses

To assess the role of *pgmA* in the growth of Mtb inside the cell, we analyzed the fitness of Δ*pgmA* by exposing different Mtb strains to host-induced stressors (Fig. 3A). After 48 hours, Δ*pgmA* exhibited approximately ∼50% susceptibility to nitrosative stress. Sensitivity levels were approximate ∼90% for oxidative stress, ∼60% for pH stress, and ∼80% for hypoxic stress. Based on the above finding we hypothesized that *pgmA* gene of Mtb must be playing a crucial role in helping Mtb survive inside the macrophages. To test this, we further estimated the relative fitness of the wild type, Δ*pgmA*, and complimented strain to replicate inside macrophages. Briefly, both resting and activated THP-1 cells were infected by giving 1:1 MOI and on the 6th day post-infection, THP-1 cells were lysed and the intracellular bacteria was assessed by CFU plating. We found that relative to the wild type and the complemented strain, Δ*pgmA* demonstrated reduced fitness in its ability to survive inside both resting and IFN-γ-activated macrophages (Fig. 3B). Consistent with the CFU assay results, we found that THP-1 macrophages infected with the Δ*pgmA* mutant strain showed increased phagosome– lysosome fusion compared to wild-type and complemented strain–infected macrophages (Fig. 3C and 3D; also see Fig. S4A). Taken together, our data highlights the central role of *pgmA* in various stress conditions, including those induced by macrophages.

**Figure 3:**
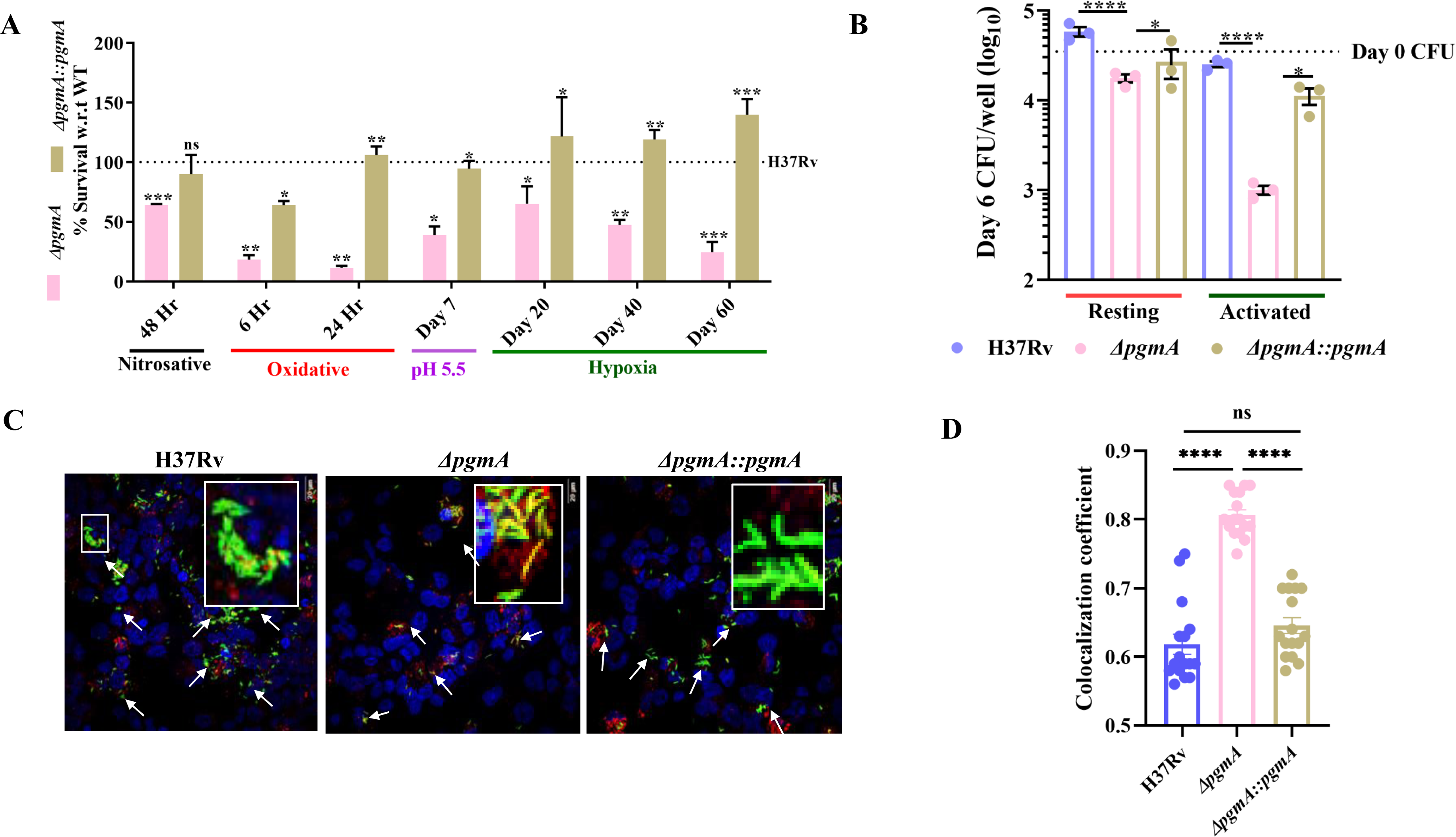
Deletion of *pgmA* renders Mtb susceptible to various stresses. (A) H37Rv, Δ*pgmA* and complemented strain was subjected to different stress conditions: nitrosative stress with 200 μM of Deta-NO for 48 h and oxidative stress with 5 mM of H_2_O_2_ treatment for 6 h. The percent survival of Δ*pgmA* relative to the wild-type strain was calculated by plating cultures at day 0 and respective time points. (B) Resting and IFNγ (10ng/ml) activated THP1 cells infected with 1:1 (MOI) of infection with H37Rv and Δ*pgmA.* In (C) and (D), the deletion of *pgmA* was observed to enhance phagosome-lysosome fusion in THP-1 macrophages infected with Mtb. In (C), THP-1 macrophages were infected with GFP-labeled strains, and images were captured using an Olympus FV3000 confocal microscope at 60× magnification (Scale bars, 20 μm). In (D), the co-localization coefficient between red and green fluorescent signals for all three strains was determined using Olympus cellSens software. The data presented for co-localisation coefficient was calculated using 16 z-stacked images obtained from two independent experiments. Statistical significance (A) to (B) and (D) was determined using unpaired, non-parametric two-tailed t-test *P < 0.05, **P < 0.005, ***P < 0.0005, and ****P ≤ 0.00005 ns = not significant. Data represent mean ± SEM (standard error mean) for technical triplicates.

### *pgmA* mediated transcriptional reprogramming help Mtb survive under nutritionally limiting condition

To further understand the role of *pgmA* in supporting the growth of Mtb during starvation, we conducted differential gene expression analysis studies to identify genes and pathways essential for *pgmA*-mediated survival during starvation. The transcriptional analysis began with an initial comparison of respective transcriptomic profiles of both H37Rv and Δ*pgmA* in starved versus normal conditions. To specifically identify genes differentially expressed by *pgmA* during starvation, we excluded the starvation-specific signal common to both H37Rv and Δ*pgmA* from our final analysis (Fig. 4A). Finally, while analyzing the final list of *pgmA*-specific genes differentially expressed during starvation revealed 26 genes to be up-regulated (red dots), while 198 genes were down-regulated (blue dots) (Fig. 4B).

**Figure 4:**
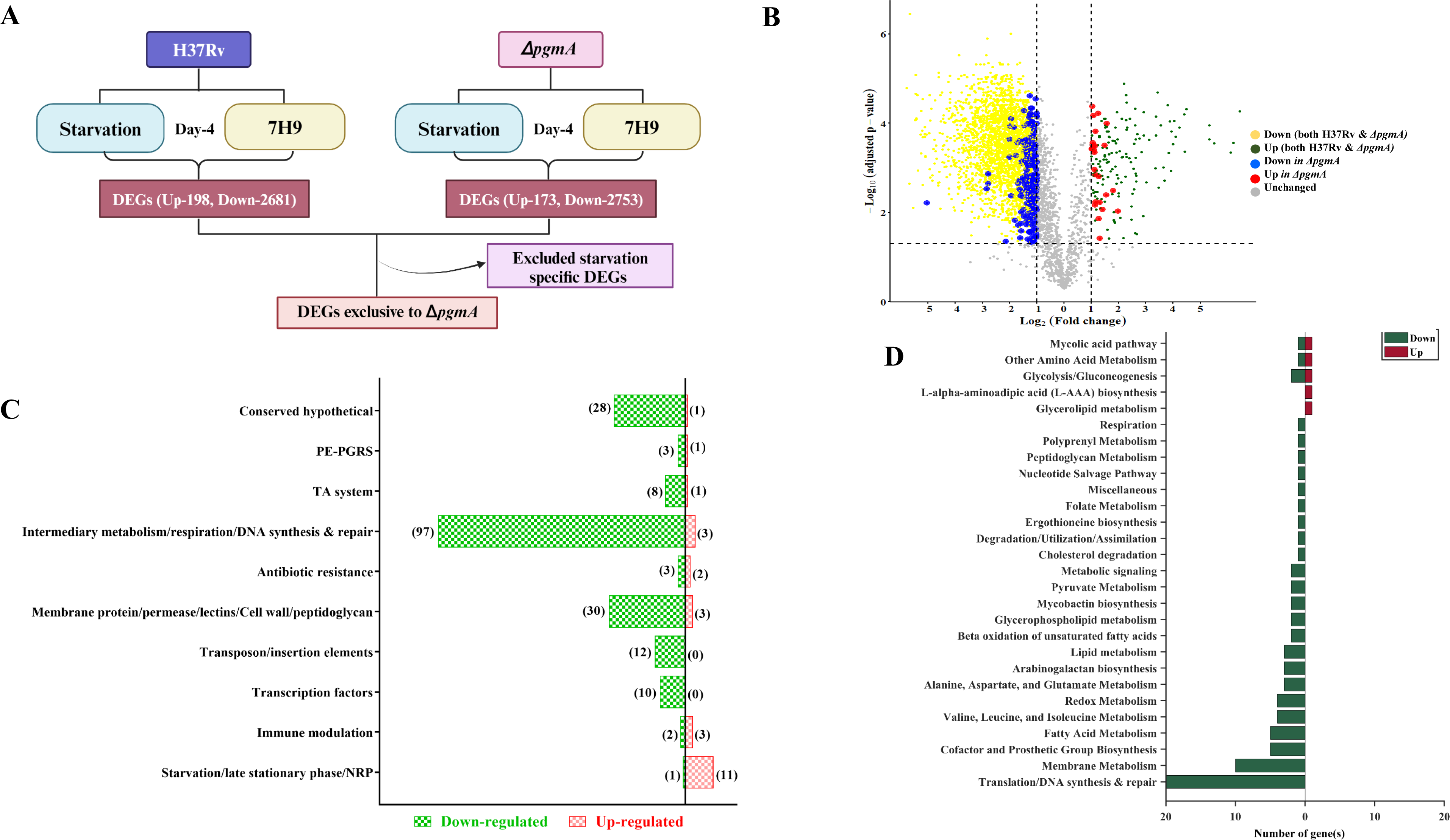
*pgmA* mediated transcriptional reprogramming help Mtb survive under nutrient limiting condition. **(A)** Schematic representation of strategy involved in RNAseq sample processing and analysis. (B) The volcano plot illustrates the results of a differential expression analysis comparing gene expression in starving versus normal conditions for both H37Rv and Δ*pgmA*. Differentially expressed genes (DEGs) were identified using a log_2_ fold change cut-off of 1 and an FDR p-value < 0.05 ^68^. The genes that are exclusively up- and down-regulated in Δ*pgmA* during starving conditions are highlighted in red and blue, respectively. Commonly up- and down-regulated DEGs in both H37Rv and Δ*pgmA* are depicted in green and yellow, respectively. In (C), Functional characterization of the uniquely regulated genes extracted using gene ontology in Δ*pgmA* is presented (Table S1). In (D), the bars represent the count of significantly altered metabolic genes involved in enriched metabolic pathways specifically for Δ*pgmA* during starving conditions. Pathway information was extracted from the genome-scale metabolic model of Mtb H37Rv, known as iEK1011 ^72^ and BioCyc genome database ^27^. Herein, the red and green colors of the bars represent the count of up- and down-regulated genes, respectively (Table S2).

Interestingly, the *Rv3290* a gene known to be a global regulator of mycobacterial persistence was found to be the most up-regulated gene of Mtb. The *pgmA*-dependent starvation gene list also included genes encoding membrane proteins such as PE-PGRS (*Rv2340c*), membrane protein (*Rv1004c*) and permeases (*Rv1999c*), metabolic enzymes like *Rv0356c* (putative thioesterase) and *Rv0223c* (aldehyde dehydrogenase) which are known to respond to oxidative stress conditions that induce membrane lipid peroxidation 25. Conversely, the list of genes that were downregulated included genes with metabolic and respiratory functions. Notable genes within this cluster include *canB* (*Rv3588c*), *pcaA* (*Rv0470c*), *hemE* (*Rv2678c*), *cyP123* (*Rv0766c*), *mbtD* (*Rv2381c*), *murG* (*Rv2153c*), *ppsD* (*Rv2934*), *glmS* (*Rv3436c*), *pfkA* (*Rv3010c*), *lipY* (*Rv3097c*), *lepA* (*Rv2404c*), *fadD2* (*Rv2948c*), *ppsB* (*Rv2932*), *gpdA2* (*Rv2982c*), *FabG5* (*Rv2766c*), *rsfB* (*Rv3687c*), *fadD22* (*Rv2948c*) and *fadD35* (*Rv2505c*). The list further included genes that encoded membrane proteins *Rv0666, Rv0488, Rv0219, Rv1382, Rv2254c, Rv0680c, Rv2307c, Rv0463, Rv0048c*, *Rv2293c, Rv0289* (*espG3*), *Rv2723, Rv2686c* (ABC transporter), *Rv2325c, Rv2403c* (*lppR*), *Rv1986, Rv2643* (*arsC*), *Rv3454, Rv0473, Rv3821, Rv1541c, Rv1217c* (ABC transporter), *Rv1146* (*MmpL13b*), *Rv1038c* (*esxJ*), *Rv1793* (*esxN*), *Rv1914c, Rv2620c, Rv0588* (*yrbE2B*), *Rv0008c*, *Rv0584* which indicates severe membrane remodelling associated with Δ*pgmA* strain under nutrient scarcity. Functional characterization of these genes using gene ontology revealed that a majority of the gene subset up-regulated belong to pathways critical for maintaining the late stationary phase, NRP (non-replicating persistence), and survival of Mtb during starvation (Fig. 4C, Table 1S). Since *pgmA* gene encodes for an enzyme that critically controls the flux of carbon flow between energy and biosynthesis (Fig. 1A), the absence of *pgmA* gene would certainly destabilize the CCM of Mtb essential for regulating optimal energy production and biosynthesis during starvation. To study this imbalance we extracted the metabolic pathways using the metabolic gene and its associated metabolic reactions from the metabolic model of Mtb iEK1011_2.0 and BioCyc genome database as described previously ^26, 27^. The bars in the graph depict the count of significantly altered metabolic genes (*x-axis*) involved in enriched metabolic pathways (*y-axis*) specifically for Δ*pgmA* during starvation (detailed in Table 2S). The coloration of the bars, red and green represents the number of up-regulated and down-regulated metabolic genes, respectively. A substantial down-regulation was observed in the majority of metabolic pathways, including 97 metabolic genes associated with membrane metabolism, fatty acid metabolism, nucleotide metabolism, and other critical pathways (Fig. 4D). This data suggests that *pgmA* modulates metabolic transcriptome in a way that helps Mtb to adapt and survive better under nutritional deprivation.

### *pgmA*-mediated regulation of carbon flux is crucial for the survival of Mtb under nutrient stress

To investigate the regulation of carbon flux in response to nutrient scarcity, we conducted a carbon flux tracing experiment using uniformly labeled U-^13^C_3_ glycerol (Gly). The aim was to discern any *pgmA*-dependent shift in the metabolic flux in Mtb exposed to nutrient-limiting conditions. To achieve this, both H37Rv and the Δ*pgmA* strains were first grown to OD: 0.6 in minimal media supplemented with labeled U-^13^C_3_ glycerol. Subsequently, to induce a starvation response, the spent media was replaced with PBST, and the strains were further cultured for 7 days. Samples were processed and LC/MS data acquisition was done at different time points (Fig. S5A). It is well-established that ^13^C_3_ glycerol is directed from the glycolytic flux through glycerate-3-phosphate ^28^. Initially, at time point zero, just before the onset of starvation, we identified a significant shift in ^13^C flux, particularly evident in trehalose and maltose-1-phosphate levels, indicating impaired G1P to G6P conversion and corresponding downstream metabolic disturbance (Fig. 5A). Following starvation, as anticipated, there was a notable impairment in glycolytic flux, as evidenced by the decreased levels of G6P and F6P, along with a reduction in the levels of intermediates in the TCA cycle in the *pgmA*-deficient strain. Conversely, trends in the biosynthetic arm, which includes trehalose and maltose-1-phosphate, remained relatively steady. A significant decrease in ^13^C labeling was also observed in the amino acid pool such as tryptophan, glutamate, alanine, proline, phenylalanine, isoleucine and tyrosine (Fig. 5B). Interestingly, the decrease in cAMP levels observed *in-vitro* calorimetric based estimation (Fig. S5B) was further confirmed by a relative decrease in the ^13^C labeling of the cAMP isolated from in Δ*pgmA* (Fig. 5B). Taken together, the above data suggest that the carbon stored in the form of glycogen under nutrient replete condition help Mtb to maintain the CCM when exposed to a stringent nutrient limiting niche inside the host (Fig. 5C).

**Figure 5:**
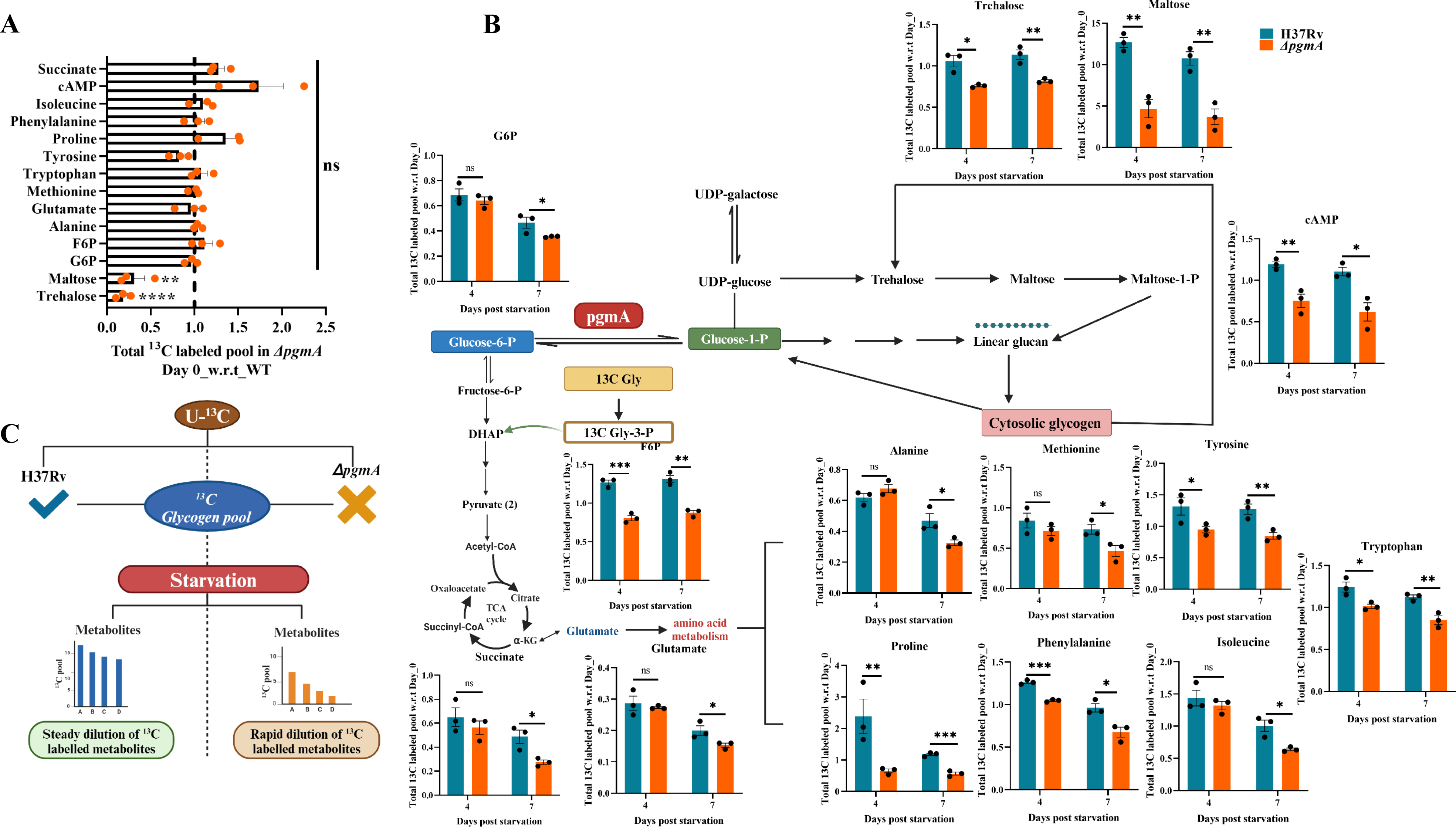
*pgmA*-mediated regulation of carbon flux is crucial for the survival of Mtb under nutrient stress. (A) The initial abundance of U-^13^C_3_-labeled metabolites in the pool prior to the onset of starvation, relative to H37RV, at time point zero. (B) Each bar for H37Rv (blue) and Δ*pgmA* (orange) represents the ratio of the total pool ^13^C_3_ labeled for individual metabolite at day 4 and 7 post starvation, this includes metabolic lineation from both G6P and G1P. (C) *pgmA* tightly regulates the carbon flux between G6P and G1P during nutrient starvation. Statistical significance between the ratio of ^13^C-labeled pools at day 4 and 7 with respect to (w.r.t) day 0 for each metabolite was analyzed using unpaired t-test *P < 0.05, **P < 0.005, ***P < 0.0005 and ****P ≤ 0.00005, ns = not significant. Abbreviations: Gly = glycerol, Gly-3-P= glycerate-3-phosphate, α-KG = α-ketoglutarate, C = carbon. Data represents mean ± SEM (standard error mean).

### *pgmA* is essential for cholesterol-specific modulation of growth and antibiotic persistence in Mtb

The role of cholesterol in the persistence and survival of mycobacteria is well-documented ^29, 30, 31^. Earlier Mtb transposon mutant library studies have predicted that the absence of *pgmA* imparts a growth advantage phenotype to Mtb in media having cholesterol as a sole carbon source (Fig. S6A) ^32^. To further confirm this finding we performed the growth kinetic studies and assessed the growth of Δ*pgmA*, wild type, and complimented strains in media harboring either glycerol or cholesterol as a sole carbon source. As previously reported, we also found that in comparison to the wild type and the complimented strain, Δ*pgmA* demonstrated an enhanced growth rate (∼3-5 fold) which was specific to media containing cholesterol as a sole carbon source (Fig. 6A). ATP being the currency of energy has a direct bearing on the growth rate of living beings. While there was no difference in the ATP levels between strains grown on glycerol, we observed that Δ*pgmA* grown on cholesterol was presented with more than 6-fold and 3-fold high levels of ATP relative to the parental and complemented strain respectively (Fig. 6B). Since it is well documented that the cAMP levels inside the pathogen are inversely correlated with the rate of growth of Mtb in cholesterol, we quantified the total cAMP levels in all three strains (Fig. 6C). While, we did not find any difference in the cAMP levels in all the three strains in glycerol, surprisingly, the cAMP levels in cholesterol media was found to be different in all three strains. While we observed a ∼3 and ∼16 fold increase in the levels of cAMP in H37Rv and the complemented strain (as compared to glycerol), surprisingly, a ∼5.7 fold decrease was observed as compared to the parent and ∼14 fold decrease was observed as compared to the glycerol media respectively in Δ*pgmA* when cholesterol was used as a carbon source. To corroborate the above findings, we conducted a comparison of the relative expression levels of certain well-known adenylate cyclase genes of Mtb in Δ*pgmA* cultured in media with glycerol and cholesterol as the exclusive carbon sources. We noticed a significant upsurge in the expression levels of all adenylate cyclases in Δ*pgmA* grown in glycerol, compared to the wild-type strain (Fig. S6B). Surprisingly, in cholesterol-grown Δ*pgmA*, there was a significant decrease in the overall expression of the same genes in relative to the wild-type strain (Fig. 6D). *prpD* gene of Mtb, which is known to be up-regulated in Mtb grown on cholesterol ^33^, was used as a control to monitor the overall expression of cholesterol induced Mtb genes.

**Figure 6:**
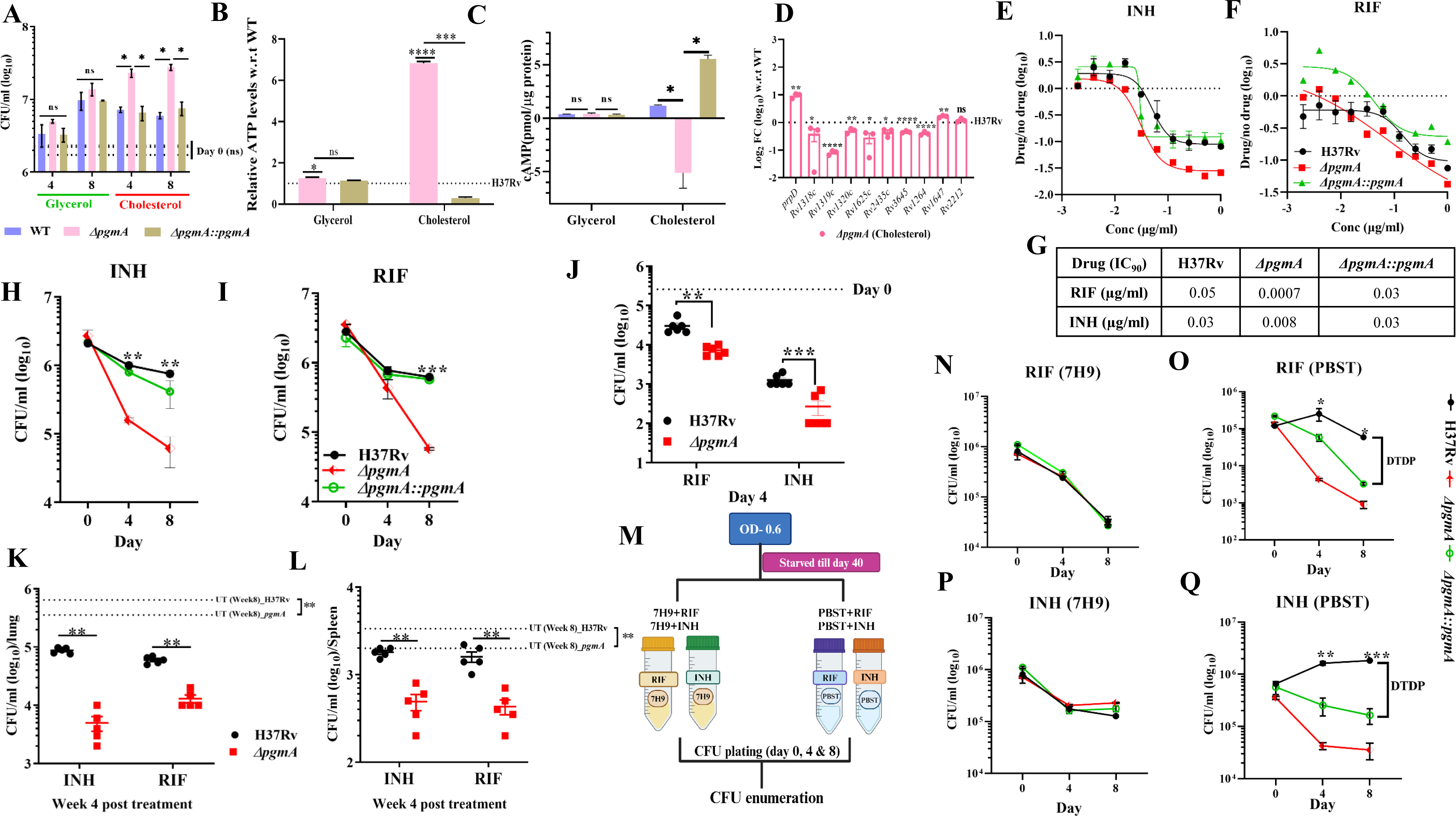
*pgmA* is essential for cholesterol-specific modulation of growth and antibiotic persistence in Mtb. **(A)** Growth curve of H37Rv and Δ*pgmA* in a minimal medium supplemented with 0.1% glycerol and 0.01% cholesterol wherein Δ*pgmA* clearly over grows in cholesterol specific media. (B) cAMP and (C) ATP measurement of H37Rv and Δ*pgmA* in 0.01%glycerol and 0.01% cholesterol. (D) The expression levels of adenylate cyclases in the Δ*pgmA* were evaluated with respect to (wrt) H37Rv under cholesterol-specific growth condition. In (E) to (G), MIC determination was done using bioluminescence bacterial strains showing cholesterol media specific susceptibility of *pgmA* to (E) INH and (F) RIF wherein, (G) showing IC_90_ values in cholesterol media. (H) RIF and (I) INH dependent time kill assay of H37Rv, Δ*pgmA* and Δ*pgmA::pgmA* in cholesterol media. (J) THP-1 macrophages were infected with H37*Rv*, Δ*pgmA*, and Δ*pgmA*::*pgmA* at a multiplicity of infection (MOI) of 1:10. After 4 h, 10x MIC of INH (0.3 µg/ml) and RIF (0.04 µg/ml) was added to the infected macrophages. At day-4 post-infection, macrophage cells were lysed, and bacillary survival was examined. CFU obtained for drug treated infected macrophages at day 4 post-infection. In (K) and (L) C57BL6 mice (*n* = 5 per group at each time point) were aerosol challenged with H37*Rv* and Δ*pgmA.* After the establishment of infection for 4 weeks, antibiotics (INH or RIF) were given through oral gavaging for next 4 weeks. A parallel group was left untreated (control). CFU enumeration was done from cell lysates of Lung **(A)** and Spleen **(B)** of Balb/c mice (*n=5*) post RIF and INH treatment, wherein, untreated group (*n=5*) was kept as a control. (M) Schematic representation showing experimental strategy for performing time kill assay on starvation induced dormant population of Mtb*. (*N) RIF and (P) INH dependent time kill assay of starvation induced dormant population in 7H9 media. (O) RIF and (Q) INH dependent time kill assay of starvation induced dormant population in PBST media. Statistical significance (A) to (D), (H) to (L) and (N) to (Q) was determined using unpaired, non-parametric two-tailed t-tests *P≤0.05 **P ≤ 0.005 and ***P ≤ 0.0005. Data represents mean ± SEM (standard error mean). Abbreviations: MOI= multiplicity of infection DTDP= drug tolerant dormant population.

The efficient regulation of CCM of any pathogen is integral to its survival and growth of the pathogen inside the host. The same holds for Mtb as well. Studies on Mtb have already established an association between mycobacterial antibiotic tolerance and CCM ^34^. However, the implications of an altered CCM in Δ*pgmA* on the pathogen’s drug susceptibility profile have never been studied. To begin with, we estimated the minimum inhibitory concentration (MIC) of H37Rv, Δ*pgmA*, and complimented strain under various growth conditions. Interestingly, we observed approximately 4-fold and 70-fold enhanced susceptibility of the Δ*pgmA* to rifampicin and isoniazid relative to the wild type and the complemented strain (Fig. 6E to 6G). Surprisingly, this phenotype was specific to cholesterol because we did not see any difference in the drug susceptibility profile within the strains under growth conditions where glycerol or an enriched media (7H9) was used as a carbon source (Fig. S6C to S6H)). Further, to study the role of the *pgmA* gene in inducing the drug tolerance phenotype in Mtb, we quantified the frequency of generation of persisters in different Mtb strains by performing a time-kill assay. Briefly, cells in the normal growth phase were subjected to INH and RIF at 10X MIC across various media specific to different carbon sources, followed by CFU plating on days 4 and 8. Similar to MIC data, we observed a cholesterol-specific decrease in the generation of persisters in Δ*pgmA* (Fig. 6H and 6I) with no observed difference in 7H9 and glycerol media (Fig. S6I to S6L). Further, we validated the above in-vitro data in both *ex-vivo* macrophage and *in-vivo* animal models. In *ex-vivo* study, THP-1 macrophages after infecting with H37Rv and Δ*pgmA* were subsequently treated with 10X MIC of RIF and INH (Fig S6M). Interestingly, in accordance with the in-vitro data, Δ*pgmA* exhibited increased susceptibility to both the first-line anti-TB drugs, underscoring the indispensability of *pgmA* in drug-induced tolerance (Fig. 6J). This data further suggests that Mtb utilizes macrophage-derived cholesterol as one of the major nutrient sources inside the cell. To corroborate our ex-vivo findings in an animal model, we administered drugs to groups of Balb/c mice infected with either the wild type or the Δ*pgmA* 4 weeks post-infection. The susceptibility of the strain to different drugs was estimated by CFU plating the lung and spleen lysates isolated from the drug-treated mice. Consistent with our findings from the macrophage study, Δ*pgmA* strain was found to be susceptible to RIF and INH (Fig. 6K and 6L). Δ*pgmA* was found to be nearly 15% and ∼10% more susceptible to INH and RIF respectively in the lungs and ∼30% and ∼40% in the spleen respectively when compared to the untreated group (Fig. S6N and S6O).

Further, it is widely recognized that mycobacteria, when subjected to nutrient deprivation, enter a state of antibiotic tolerance or resistance ^35, 36^, a phenomenon primarily attributed by number of factors such as membrane lipids like PDIMs, toxin-antitoxin systems and transcription factors ^37, 38^. To investigate if *pgmA* gene of Mtb contributes to this phenotype, we starved all three strains of Mtb for 40 days followed by treating each of these strains by RIF and INH at again concentration of 10X MIC under both nutrient replete (7H9) and nutrient depleted conditions (PBST) (Fig. 6M). Interestingly, the data from the time-kill assay revealed that starved cultures of the wild type, Δ*pgmA* and the complimented strain when treated with both the drugs under nutrient replete condition generated similar kill kinetics suggesting generation of equal number of persisters (Fig. 6N and 6P). On the contrary, the starved wild type strain was found to be completely tolerant to both the tested drugs. Intriguingly, this starvation induced drug tolerance phenotype was completely dependent on the presence of *pgmA* gene since absence of this gene in Δ*pgmA* obliterated the generation of drug tolerant bacteria as evident in the time-kill assay (Fig. 6O and 6Q). Overall, the data suggests that *pgmA* regulated CCM modulates drug susceptibility and is critical for inducing drug tolerance phenotype under nutrient limiting condition typically encountered by the pathogen inside the host.

### Deletion of *pgmA* impairs growth of Mtb in mice

Finally, to elucidate the role of *pgmA* in mycobacterial virulence, we infected groups of C57BL/6 mice with wild-type, Δ*pgmA*, and the complemented strain, and assessed virulence by measuring bacillary load via CFU plating at weeks 4, and 8 post-infection. A significant reduction in bacillary load was observed in the lungs and spleen of animals infected with Δ*pgmA* compared to those infected with H37Rv and the complemented strain. The reduction was more pronounced in spleen. Additionally, upon gross examination, lungs and spleens isolated from Δ*pgmA*-infected mice exhibited reduced inflammation and fewer visible granulomas on the surface compared to organs from wild-type-infected mice (Fig. 7C). Histopathological analysis further demonstrated that the lungs of mice infected with Δ*pgmA* exhibited diminished disease pathology and a lower granuloma score relative to those infected with wild-type Mtb (Fig. 7D and 7E). Overall, these findings suggest that *pgmA* plays a crucial role in modulating Mtb virulence and its absence appears to compromise the fitness and ability of Mtb to proliferate within the host.

**Figure 7:**
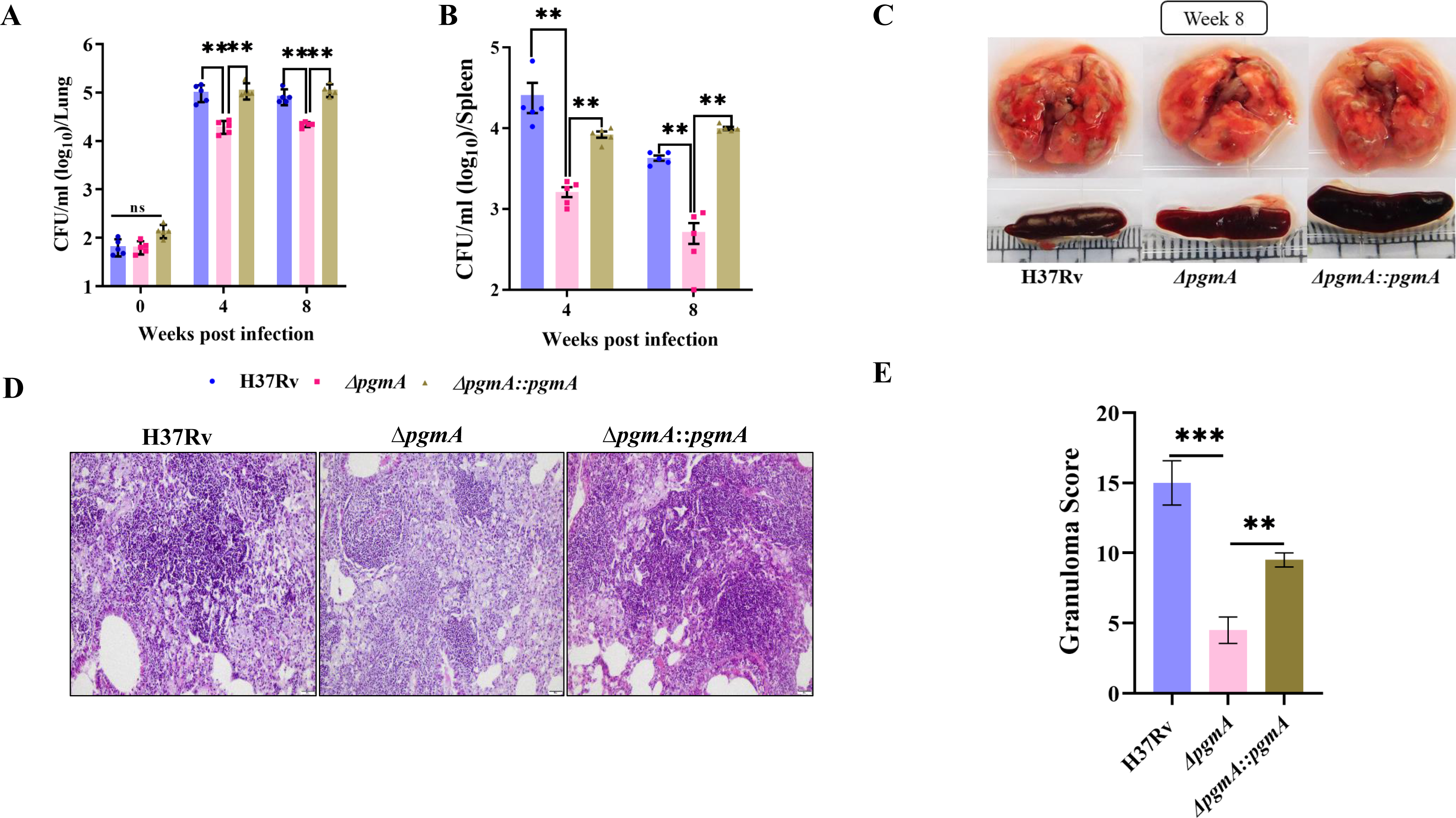
Deletion of *pgmA* impairs growth of Mtb in mice. Female C57BL6 mice (n=5) were subjected to aerosol challenge with *H37Rv, ΔpgmA, and ΔpgmA::pgmA* strains to evaluate disease persistence in (A) lungs and (B) spleen at specified time intervals. (C) Gross pathology of the infected lungs and spleen was assessed after 8 weeks post-infection. Lung sections stained with hematoxylin and eosin were imaged at 20x magnification and analyzed for Mtb-induced histopathological changes. The alteration in lung morphology, including granuloma formation (E), is depicted. Statistical significance was determined using unpaired, non-parametric Mann-Whitney U test in (A) and (B) **P ≤ 0.08, and unpaired two-tailed t-test in (E) *P≤0.05 **P ≤ 0.005 and ***P ≤ 0.0005. Data represents mean ± SEM (standard error mean).

**Figure 8:**
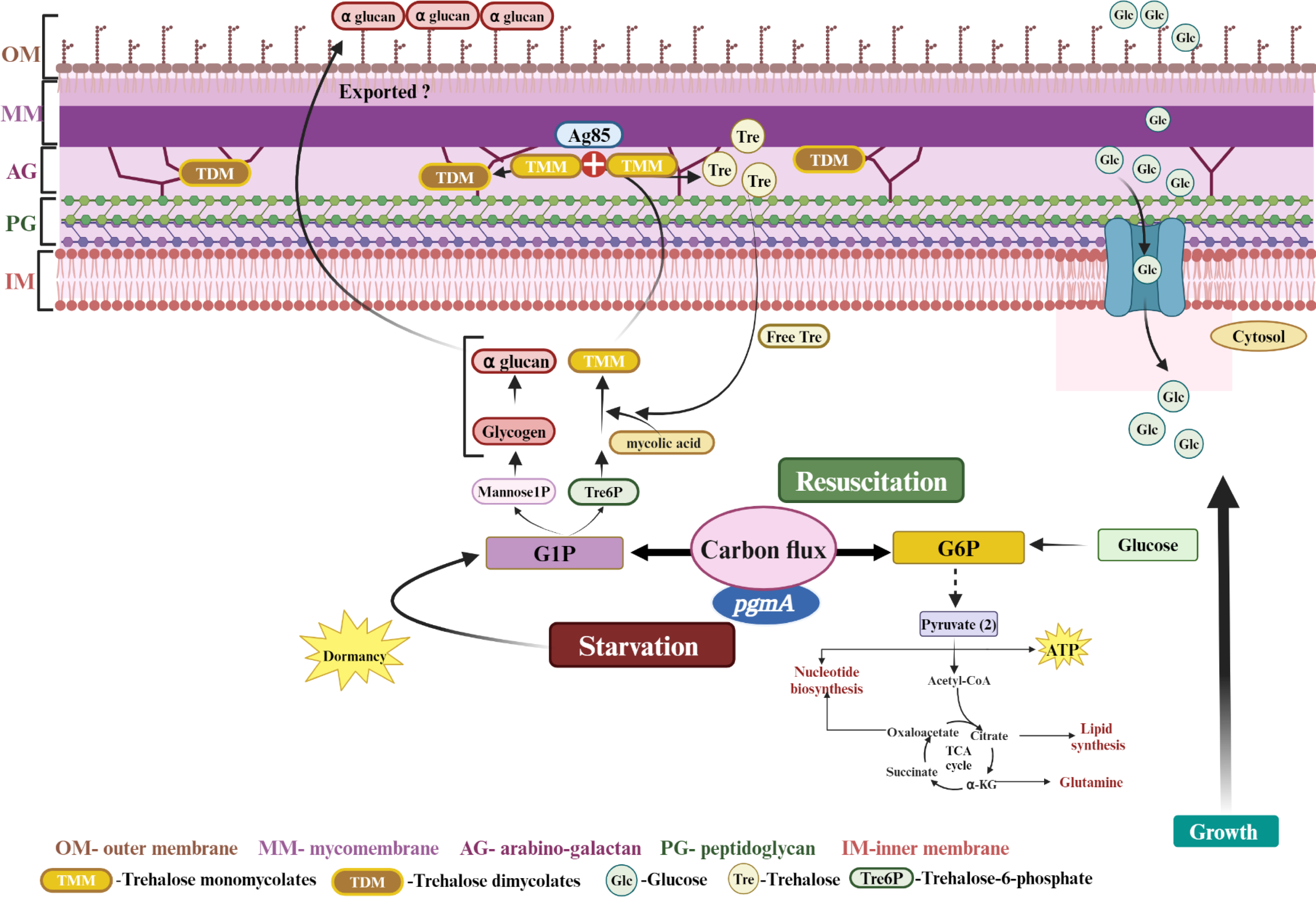
Schematic representation of the pivotal role played by the *pgmA* in regulating carbon flux. *pgmA* orchestrates the conversion of G6P to G1P, thereby contributing significantly to stabilizing the CCM of Mtb. This regulation directs carbon flux towards either energy production or biosynthesis, contingent upon diverse growth conditions. Consequently, Mtb can thrive by tapping into stored carbon reservoirs during periods of nutrient scarcity and during resuscitation phases. Moreover, the *pgmA*-mediated production of G1P acts as a critical precursor for metabolites essential in synthesizing membrane-associated lipids, pivotal for maintaining cell wall integrity. Hence, the *pgmA*-dependent carbon flux switch assumes indispensable importance for Mtb throughout disease progression and antibiotic tolerance. This flux facilitates adaptation to varying environmental conditions and ensures metabolic adaptability crucial for survival and virulence.

## Discussion

The phenotypic plasticity of Mtb is defined by its extraordinary ability to adapt to a variety of adverse environmental changes faced within the host. Although a stable central carbon metabolism ensures a continuous flow of energy and biomolecules essential for maintaining critical cellular processes, adaptability is defined by the ability of the pathogen to differentially regulate the flux of the available nutrient between these two pathways to maximize the efficiency of utilization of the available resources. Our data suggest that conversion of G6P to G1P by *pgmA* is one such step that Mtb modulates to regulate the flux of carbon under nutrient limiting and reactivation conditions ^39^. In this study, we found that the *pgmA*-mediated conversion of G6P to G1P or vice versa helps stabilize the CCM of Mtb by directing the carbon flux towards energy or biosynthesis depending on the growth condition. Interestingly, data from the growth and ^13^C carbon flux analysis suggests that glycogen acts as a “carbon capacitor” and helps Mtb to survive by releasing the stored carbon under nutrient-limiting conditions and during reactivation. Our lipid data reveals that the *pgmA*-dependent generation of G1P acts as a precursor for the metabolites essential for the synthesis and transportation of membrane-associated lipids critical for maintaining the cell wall integrity and Mtb virulence ^40, 41^. The reactivation kinetics clearly demonstrates the importance of this flux in allowing Mtb to regulate the flow of carbon between different catabolic and biosynthetic pathways as and when needed. Additionally, we report that *pgmA*-dependent growth modulation in cholesterol drives the generation of antibiotic persistence during tuberculosis infection. Finally, our study demonstrated that the *pgmA* gene of Mtb is indispensable for the long-term growth and survival of the pathogen inside the host.

Similar to our findings, using a transposon mutant library, it was recently reported that the *pgmA* gene was essential for Mtb to grow under both starvation and reactivation conditions ^42^. This phenotype is not restricted to mycobacteria and has been reported in several other bacterial pathogens like *E. coli, S. typhimurium*, and *Vibrio* spp, wherein mutants with impaired glycogen metabolism compromise fitness during carbon starvation ^43, 44^. Failure of *pgmA* deficient mutant to divert the carbon flux into synthesis of critical macromolecules essential for the generation of membrane-associated lipids, compromises the cellular morphology and membrane integrity, rendering the mutant strain susceptible to exposure to multiple stress conditions. The reported down-regulation of *pgmA* under hypoxia condition is very intriguing ^45^. This together with our findings suggests that, despite showing a decrease in the transcript levels, *pgmA* is essential for the growth of bacteria under hypoxic conditions. Furthermore, our findings indicate that Phosphoglucomutase-β (*Rv3400*), the protein of which has recently been studied ^46^, did not show redundancy upon removal of *pgmA*.

Defects in the formation of biofilm in Δ*pgmA* can be attributed to its inability to synthesize and transport TDM also referred to as cording factor ^47, 48^. The importance of a balanced and efficient regulation CCM has been reported to be essential for the synthesis of mycolic acid critical for cell wall integrity and biofilm formation in Mtb ^49, 50, 51^. Since mycolic acid are tightly anchored to the arabinogalactans (AGs), the inability of the Δ*pgmA* to synthesize common UDP-linked sugar precursors such as trehalose, essential for the synthesis of AGs, could also be one of the reasons for the absence of surface mycolic acid. The decrease in the levels of apolar membrane lipids like TAGs and a fraction of PDIMs from the cell membrane was very intriguing. Lipid separation was performed during the late log and early stationary phases of cells, at which point it is know that Mtb switches to stored nutrients. TAG serves as a reliable long-term energy source with a lower molecular mass than glycogen ^52^. Its consumption in *pgmA* lacking strain indicates reliance on TAG in the absence of glycogen. Interestingly, the decrease in PDIM levels in Δ*pgmA,* correlates with our RNAseq data where we observed a Δ*pgmA*-specific down-regulation of *ppsD* (*Rv2934*) and *ppsB* (*Rv2932*) genes essential for the biosynthesis of PDIM in Mtb. Consistent with recent reports ^53^, we observed increased sensitivity of Δ*pgmA* to the peptidoglycan-targeting drug vancomycin. This finding supports the hypothesis that the *pgmA* gene in Mtb is essential for the synthesis and integrity of its glycolipid-rich cell wall. The difference in the overall morphology, more specifically the difference in the size, observed in Δ*pgmA* we believe is a strategy adopted by the Δ*pgmA* to remodel the cell wall to compensate for the lack of membrane glycolipids. The possibility of this being part of a strategy to conserve energy by the Δ*pgmA* cannot be ruled out. Similar observations have been reported in other bacterial species including *E. coli*, *Vibrio spp.*, *S. aureus*, and *M. luteus*, where cytoplasmic shrinkage occurs during dormancy to conserve energy ^44, 54, 55^. Alternatively, failure of the Δ*pgmA* to synthesize and store glycogen limits the ability of the mutant to generate ATP under nutrient-limiting conditions. RNAseq and ^13^C carbon flux data provide clear evidence of shifting of major metabolic pathways in the absence of *pgmA*. Needless to say, this shift clearly blunts the ability of the pathogen to replicate and survive inside the host for an extended period. This can be explained by comparing the growth kinetics of Mtb and Δ*pgmA* in this study. The indispensability of *pgmA* throughout the disease process underscores the requirement of glycogen as a carbon capacitor in maintaining both the actively replicating and persistence phase of the infection in tuberculosis.

The *pgmA*-dependent growth modulation of Mtb in cholesterol was very intriguing. Although, the cAMP-dependent growth modulation of Mtb in cholesterol is already known ^56, 57, 58, 59^, we for the first time implicate *pgmA* gene of Mtb in modulating the cAMP levels in the presence of cholesterol. Furthermore, the role of cAMP in mediating the drug susceptibility of Mtb has emerged as a recent focal point of research, yielding some compelling findings ^58, 60, 61, 62^. The cholesterol-specific imbalance in carbon flux in the absence of *pgmA* may have contributed to the decrease in cAMP and increased ATP levels, potentially explaining the overgrowth phenotype observed during the growth in cholesterol-specific conditions. Moreover, the difference in growth rate also altered the drug susceptibility profile of Δ*pgmA* on cholesterol. Interestingly, the observed drug susceptibility signature of Δ*pgmA* under *ex-vivo* and *in-vivo* growth conditions were identical to the data obtained under *in-vitro* cholesterol growth conditions. This underscores the fact that host-derived cholesterol is one of the major sources of nutrients available for Mtb to survive inside the host. The *pgmA*-dependent differences observed in the drug susceptibility profile were a consequence of the increase in the growth rate or due to the differences observed in the cAMP levels is not clear and requires further studies. Overall, a detailed mechanistic understanding of *pgmA*-mediated cAMP-dependent growth modulation and its implications on drug tolerance will be an interesting area for future research. Further, persisters are recognized for their resistance to most drugs ^15, 35^. In our experiment, we noticed that, in Mtb lacking *pgmA*, starvation-induced drug tolerance was completely abrogated and the strain was found to be more susceptible to drugs under non-permissive nutritional conditions. The data suggests a dominant role of *pgmA* in modulating drug susceptibility phenotype of Mtb under nutrient-limiting conditions. Surprisingly, the above phenotype was very specific to the nutrient limiting conditions emphasizing the importance of the *pgmA*-regulated carbon flux in inducing drug tolerance phenotype in Mtb. Finally, the indispensability of *pgmA* throughout the disease process underscores the essentiality of spacio-temporal regulation of the carbon flux in both the actively replicating and persistence phase of the infection in tuberculosis. This balance was found to be crucial which otherwise caused accelerated clearance of Δ*pgmA*, whether due to drug-induced processes within the host or its incapacity to persist *in-vivo*. Also, the role of membrane lipids have been reported to play a critical role in its ability to survive within macrophages and to maintain a successful infection ^42, 63, 64, 65^. Overall, these observations underscore the role of *pgmA* in both antibiotic tolerance and disease persistence.

Based on our earlier and current findings we propose that the *pgmA*-driven reversible conversion of G6P to G1P is a critical step that regulates the central carbon metabolism of Mtb both during active growth and under nutrient-deprived conditions inside the host. Our data also highlighted the role of *pgmA* in both maintaining the integrity of the cell wall and cAMP-dependent modulation of the growth of Mtb inside the host. More importantly, the role of *pgmA* in modulating the drug susceptibility phenotype of Mtb inside the host has never been previously reported. In conclusion, our study identified a key enzyme that not only regulates the central carbon metabolism but also helps impart drug tolerance phenotype to Mtb inside the host. Our findings strongly suggest that inhibitors of *pgmA* as an adjunct might potentiate the existing regimen against tuberculosis.

## Materials and Method

### Bacterial strains, plasmids and culture conditions

All the experiments were done using *the Mycobacterium tuberculosis H37Rv* strain. The Δ*pgmA* (*ΔRv3068c*) strain as annotated in Mycobowser was generated by replacing the *Rv3068c* gene with a hygromycin resistance cassette using the pJM1 suicide vector (Figure S1). To study gene-specific effects, a complementation strain was generated by unmarking the deletion mutant. This whole cassette was excised using Cre recombinase. The unmarked mutant strain was then transformed with pMV261 harboring the *pgmA* gene with a constitutive mycobacterial *Phsp60* promoter. GFP-expressing strains were prepared by electroporating pMV261: eGFP plasmid followed by selection on hygromycin containing 7H11 plates with green colonies indicative of plasmid-bearing bacilli. The cultures were grown in Middlebrook 7H9 broth (Difco) supplemented with 0.05% Tween-80 (sigma) and 10% OADS (albumin, glucose, NaCl, oleic acid), 0.2% glycerol (sigma) enrichment at 37^0^C, 100 rpm. Colony-forming unit (CFU) plating was done on 7H11 media supplemented with 10% OADS and 0.5% glycerol. Hygromycin and Kanamycin were used at concentrations of 50µg/ml and 25µg/ml respectively. Carbon source-specific experiments were done in minimal media (0.5 g/litre asparagine, 1 g/litre KH2PO4, 2.5 g/litre Na2HPO4, 50 mg/litre ferric ammonium citrate, 0.5 g/litre MgSO4⋅7H2O, 0.5 mg/litre CaCl2, 0.1 mg/litre ZnSO4) containing 0.1% (vol/vol) glycerol and 0.01% (wt/vol) cholesterol. For biofilm formation, Sauton’s media was used and prepared using 0.5 g of KH2PO4, 0.5 g of MgSO4, 4 g of L-Asparagine, 2 g of Citric acid, 0.05 g of Ferric Ammonium Citrate, 60 mL of glycerol in 900 mL of water followed by adjusting the pH to 7.0 with NaOH. The MIC determination was conducted utilizing a bioluminescence-based plasmid, pMV306hspLux+G13 (Addgene#26161). The plasmid was introduced into competent cells prepared with 10% glycerol for each strain via electroporation. Subsequently, the transformed cells were plated onto hygromycin and zeocin 7H11 plates, with concentrations of 50µg/ml and 25µg/ml, respectively.

### Glycogen estimation assay

The glycogen estimation kit measures glycogen enzymatically by detecting free glucose. Therefore, the estimation was done by starving bacterial cells in PBST to avoid the assay’s limitation of background free glucose interference present in 7H9. The estimation was done at day 2, 4 and 7 (Fig S2A). To evaluate glycogen levels, cells were disrupted using zirconia beads in a solution comprising 25 mM citrate at pH 4.2 and 2.5 g/liter sodium fluoride, all maintained at 4°C. The resulting mixture was clarified through centrifugation at 14,000 g for 10 minutes, and the resulting supernatant was filtered and utilized for glycogen measurement following the guidelines provided by the EnzyChrom glycogen assay kit (Bioassay Systems), as per the manufacturer’s instructions.

### EtBr permeability assay

The cell wall permeability was assessed by EtBr dye and was adapted as originally mentioned earlier ^66^. Concisely, cells were harvested in a 7H9 complete medium without detergent until A_600_ of 0.6-0.8. Following a PBS wash, cells were introduced into wells of a black clear-bottom 96-well plate containing EtBr solution prepared in PBS at final concentrations of 2 µg/ml. Fluorescence readings were recorded every minute for 60 mins at 530/590 nm. PBS was kept as a blank and the readings obtained were minus with the blank.

### Lipid extraction and thin-layer chromatography

Cultures were grown in 10 ml 7H9 media till the late log phase (A_600_ ∼1 to 1.5). The cells were then washed with PBS and re-suspended in chloroform-methanol (2:1, vol/vol) mix for overnight at 37^0^C and 100 rpm. This was followed by sequential extraction with chloroform-methanol (1:1, vol/vol) and chloroform-methanol (1:2, vol/vol). For analysing lipid defects in biofilms, cells from biofilm were taken and washed with PBS followed by sequential chloroform-methanol treatment. For lipid analysis in TLC, 10 μl of each lipid was spotted onto a pre-dried TLC plate (Silica gel; Sigma) at a distance of 2 cm upward from the end of the plate. The solvent systems used for lipid analysis are listed in Table 1S.

### Transmission electron microscopy

Specimens for TEM analysis underwent processing according to established procedures outlined elsewhere ^67^. To summarize, cells were cultured in 7H9 broth (lacking Tween 80), subsequently starved in PBST for 7 days, and approximately 5 × 10^7^ cells were harvested by centrifugation. The collected cells were then fixed using a combination of 2.5% paraformaldehyde and 2.5% glutaraldehyde. TEM was carried out utilizing a JEM1400 Flash TEM operating at 120 kV (ATPC, Regional Center for Biotechnology, India), equipped with tungsten filament as electron source and highly-sensitive sCMOS camera. The cell size was measured for *n=50* cells for each strain and the analysis was done using RADIUS 2.0 EMSIS (build 14530) inbuilt software.

### Biofilm formation

Biofilms of each strain were cultivated in 6-well plates by introducing 3 ml of Sauton’s medium (excluding Tween-80 or tyloxapol) having 4% of fully saturated planktonic culture. These dishes were then placed in a stationary incubator at 37°C, under humid conditions with 5% CO_2_, undisturbed for 5 weeks.

### *In-vitro* stress assay

To decipher susceptibility to starvation and pH stresses, A_600_∼0.5-0.6 mycobacterial cultures were washed with phosphate-buffered saline (PBS) containing 0.01% Tyloxapol (PBST), and a single-cell suspension was inoculated at an A_600_ of ∼0.5 in PBST and ∼0.2 in 7H9-ADC (pH 4.5). To assess viability after hypoxic stress, bacterial strains were inoculated with a headspace of 15% at an A_600_ of ∼0.1 in 7H9 medium containing 1.5 μg/ml methylene blue (colorimetric redox indicator of dissolved oxygen) in vacutainer vials at 37°C. For nitrosative and oxidative stresses, 200 µM DETA-NO and 50 µM CHP were used.

### Confocal laser microscopy

The eGFP-expressing strains were used for confocal microscopy experiments. THP-1 macrophages were seeded onto glass coverslips in a 24-well plate at a density of 5x10^5^ cells per well and were pre-activated with 10 ng/ml of human IFN-γ (Peprotech, USA) for 24 hours prior to infection. Bacterial cultures were cultivated until they reach the early log phase (A_600_∼0.6-0.7), subsequently harvested and then rinsed with fresh RPMI-1640 media. Following this, a single-cell suspension was prepared using the gentle spin method (centrifugation at 120 g for 10 minutes). The activated THP-1 cells were infected with a multiplicity of infection (MOI) of 1:10. After a 4-hour incubation period, the cells were washed three times with PBS to eliminate extracellular bacteria, and the media (RPMI supplemented with 10% FBS) was replenished. Subsequently, 24 hours post-infection, the cells were treated with 50nM LysoTracker Red DND-99 (Invitrogen Life Technologies, CA, USA) in complete RPMI media for 30 minutes at 37°C with 5% CO_2_ and then fixed with 4% paraformaldehyde in PBS. Coverslips were mounted with ProLong Diamond antifade with DAPI (Molecular Probes by Life Technologies, CA, USA). Finally, the slides were analyzed using an Olympus Confocal Laser Scanning Microscope (CLSM). The presence of GFP-expressing mycobacteria co-localizing with LysoTracker Red was determined by examining approximately 100 phagosomes. The CFU enumeration of all three strains was also determined by infecting activated and non-activated THP-1 macrophages with an MOI of 1:1 on day 6.

### Genome wide expression analysis

Mtb cultures, conducted in triplicate, were harvested in 7H9 media until reaching OD-0.6. Simultaneously, cultures from each strain (in triplicate) were starved (initial OD-0.5) until day 4. Subsequently, cultures from both media conditions were resuspended in Trizol. The pelleted Trizol mix underwent bead beating (using 0.2 µM zirconia beads) five times for 40 seconds each, with 1-minute intervals on ice to cool the sample. RNA was then isolated using the chloroform and isopropanol method. The total RNA was outsourced for subsequent DNAase treatment and transcriptomic analysis. Raw reads were filtered using Trimmomatic to eliminate low-quality scores and adapters. The filtered reads were aligned to the Mtb reference genome (https://www.ncbi.nlm.nih.gov/datasets/taxonomy/1773/) using splice-aware aligners such as HISAT2 to quantify reads mapped to each transcript. The alignment percentage of reads ranged between 97.9% and 99.16% for all samples. The total number of uniquely mapped reads was counted using feature counts. The uniquely mapped reads were then subjected to differential gene expression analysis using DeSeq2. Differential expression analysis of genes was performed for the starving condition versus the normal condition for both the *pgmA* mutant (*ΔRv3068c*) and H37Rv strains of Mtb, respectively, using a two-sample t-test. Genes with fold-changes < 0.5 or > 2 ^68^ and false discovery rates (FDR) < 0.05 were considered differentially expressed genes (DEGs) in the starving condition for both Δ*pgmA* and H37Rv respectively.

### Liquid Chromatography–Mass Spectrometry/Metabolomics

Mtb cells were first harvested till OD-0.6 in Uniformly labeled (U)-13C3:12C glycerol at a concentration of 0.1% with 13C //12C, ¼: v/v. Further, cells grown in 25% labeled glycerol were then transferred to starved media (PBST) with an initial OD of 0.5 till days 4 and 7. The starved cultures of mutant and parental strains were then washed twice with chilled 1X PBS. Bacterial cells were then lysed in extraction buffer (acetonitrile: methanol: water; 40:40:20) using 0.1mm zirconia beads in a bead beater (5 cycles/40s pulse each). After each cycle, the cells were kept on ice. The lysed cells were pelleted by centrifugation at 5000rpm for 10 minutes at 4°C, and the supernatant was filtered twice using a 0.2µ filter. The supernatant was dried using a speed vacuum at room temperature and stored at -80 till further analysis. Finally, for sample acquisition, the samples were resuspended in an equal ratio of methanol and water. The data acquisition was done on an orbitrap fusion mass spectrometer (Thermo Fisher Scientific) coupled with a heated electron ion source. The acquisition method has been followed as per the published protocols ^69, 70^ with minor modifications. The MS1 mass resolution was kept at 120,000 and for MS2 60,000. Separation of the extracted metabolites was done on UHPLC Dionex Ultimate 3000. The data was acquired in both the reverse and HILIC phases in both positive and negative ionization modes. The reverse phase column was HSS T3 and the HILIC column was XBridge BEH Amide (Water Corporation). For the reverse phase, the mobile phase consists of solvent A (water +0.1% formic acid) and solvent B (methanol + 0.1% formic acid). The elution gradient started with 1% B to 99% A over 10 min with a flow rate of 0.3 ml/min. For the HILIC, the mobile phase consists of solvent A (20 mM ammonium acetate in water) at pH 9.0 and solvent B (100% acetonitrile). The elution gradient started with 85% B to 15% A over 14 min with a flow rate of 0.2 ml/min. The injection volume was 5ul. Quality control (QC) sample was run between the samples to monitor the signal variation and RT shift. Data were analyzed using EL-MAVEN (v0.12.1-beta) software and ions were identified based on mass accuracy within 10 parts per million and 1 min retention time cut-off based on the in-house library of the metabolites of interest. The isotope natural abundance was corrected using AccuCor ^71^.To analyze isotopologues, the ratio of metabolite isotopic labeling was calculated by dividing the peak area ion intensities of total ion intensity of all labeled species of each metabolite w.r.t the parental strain.

### ATP estimation

ATP levels were determined by pelleting approximately 10^8^ cells at 4000 rpm. These cells were then washed twice with PBS and subsequently resuspended in 1 ml of PBS. Next, the cells were lysed using 0.22 µM zirconia beads in a mini-bead beater. Following this, the supernatant was collected after centrifuging the cell suspension at 14000 rpm. The supernatant was further processed by heating it to 95°C and subsequently filtered through 0.2µ filters. The resulting cell lysate was then subjected to ATP estimation, following the manufacturer’s protocol, utilizing the BacTiter-Glo™ Microbial Cell Viability Assay kit from Promega. Luminescence readouts were recorded and subsequently normalized per µg of protein.

### cAMP determination

To assess the cAMP levels within cells, approximately 10^8^ cell suspensions were subjected to centrifugation at 4000 rpm. The resulting bacterial pellet was reconstituted in 1 ml of 0.1 M HCl, and the cells were disrupted using 0.1 mm zirconia beads in a mini Bead Beater from BioSpec Products, USA. Subsequently, beads and bacterial remnants were separated through centrifugation, and the resulting supernatants were employed for cAMP quantification using the manufacturer protocol provided in a direct immunoassay kit by Enzo (ADI-900-067A). The absorbance was taken at 405 nm and the data was normalized per µg of protein.

### RT-qPCR

Total RNA was isolated from the respective bacterial strains under specific conditions using Trizol (TaKaRa) following the manufacturer’s protocol. In brief, bacterial cells were washed with PBS and resuspended in Trizol. The cells were lysed by bead beating using 0.22 µ sized zirconia beads (5 cycles /with intermittent cooling steps of 1 minute each). Subsequently, RNA was extracted and precipitated using chloroform-isopropanol treatments. The resulting total RNA was treated with Turbo DNaseI (Ambion) to remove DNA contamination. 3000µg equivalent of DNase-treated RNA was used to generate cDNA using SuperScript IV Reverse Transcriptase (Invitrogen). Finally, 1µg equivalent of cDNA was utilized to set up the qPCR reaction with SYBR Premix Ex Taq by Takara Bio on QuantStudio 6 Flex. *sigA* was used as the internal control and the ΔΔCt method was applied to determine the relative gene expressions concerning specified control conditions. *prpD* (*Rv1130*) was included as a positive control for cholesterol utilization.

### MIC and antibiotic tolerance experiments

The MIC was determined using bioluminescent bacterial strains with an OD of 0.5 (mid-log phase) that were washed with PBS. MIC values were determined in 7H9, glycerol, and cholesterol media. A 96-well plate was used for serially diluting the drugs, starting with twice the maximum concentration of the drug and media in the first well. Cells were added at a final density of OD 0.01 to the wells containing drugs and incubated at 37°C for 3 days. Luminescence was then measured and dose-response curves were generated to calculate IC90 values. Time-kill kinetics to assess *in-vitro* antibiotic susceptibility was conducted independently in 7H9, glycerol, and cholesterol-specific media. Mid-log-phase cultures were collected and transferred into their respective media until they reached an A_600_ of approximately 0.2. 10X MIC drug concentration was then added to these cultures, and CFU enumeration was performed on days 0, 4, and 8. For antibiotic susceptibility testing in THP-1 cells, the macrophages were infected at MOI of 1:5. After a 4-hr infection period, extracellular bacteria were removed through successive PBS washes. RPMI1640 media containing a drug concentration of 10X MIC was then added to each well. Additionally, an untreated control group was maintained for each drug. CFU enumeration of intracellular bacterial load was monitored by cell lysis by PBST having 0.01% triton-100 on day 4.

### Infection experiment in mice

For the pathogenesis study, C57BL/6 mice, aged 4 to 6 weeks, were housed in individually ventilated cages at the in-house BSL-3 facility. The mice were cared for in strict accordance with established animal ethics and guidelines. For the aerosol infection, groups of mice were exposed to a controlled bacilli concentration of 80-150 bacilli per mouse using the inhalation exposure machine, Glas col, LLC. At 24 hours post-exposure, five mice per strain were sacrificed to assess bacterial deposition. To measure bacterial loads in the lungs and spleens at various specified time points, organs were aseptically collected and homogenized in 2 ml of sterile normal saline. Subsequently, the homogenates were serially diluted and plated in triplicate on 7H11 agar with 10% OADS, supplemented with polymyxin B (50 mg/ml), amphotericin B (10 mg/ml), trimethoprim (10 mg/ml), vancomycin (5 mg/ml), carbenicillin (100 mg/ml) and cycloheximide (5 mg/ml) to prevent contamination and assess the bacillary load. For the in-vivo antibiotic tolerance study, BALB/c mice, aged 4-6 weeks were used. For the establishment of infection, animals were kept under care for 4 weeks. After 4 weeks, n=5 mice were sacrificed for bacillary load enumeration and consequently, mice were randomly grouped and subjected to treatment with 10 mg/kg INH and RIF, respectively through oral gavaging 5 days/week for 4 weeks. A parallel group was maintained as an untreated control. Bacillary survival was evaluated by sacrificing mice at 0, 4, and 8 weeks as mentioned earlier.

### Statistical analysis

Each *in-vitro* and *ex-vivo* CFU experiments were repeated thrice independently. The statistical significance was assessed using the student’s t-test in all the *in-vitro* and *ex-vivo* experiments and the Mann Whitney U-test in animal experiments. Statistically significant results were defined as having values P <0.05, P <0.005, and P <0.0005. All the analyses for RNA Seq were performed on MATLAB and R software. PCA was performed using the “prcomp” package of R. The “ttest2” function in MATLAB was applied to perform the two-sample t-test. PCA plot, volcano plot, and heatmap were generated in R using ggplot2, gplots, and Complex Heatmap packages.

### Ethical statement

The animal experimentation procedure received approval from the Animal Ethics Committee at THSTI, Faridabad, India (IAEC approval numbers 166 and 211). This approval was granted following the guidelines established by the Committee for Control and Supervision of Experiments on Animals (CCSEA), an entity under the Government of India.

## Acknowledgment

**Funding:** We acknowledge the funding from Indian Council of Medical Research, India (Project ID: EM/Dev/SG/212/7864/2023) to A.K.P. and from the Translational Health Science and Technology Institute (THSTI) core grant to A.K.P. and S.C. TS was supported by Council of Scientific and Industrial Research (File no. 09/1049(0036)/2019-EMR-I) PhD fellowship. The sponsors had no involvement in the planning, data collection, analysis, or interpretation, in the preparation of the report. **Facility:** We would like to acknowledge the support provided by the Bio-safety level 3 (BSL3), MultiOMICS, Histopathology and the Experimental Animal Facility (EAF) at THSTI. **Software:** The graphical abstract depicting the findings was created using Biorender Software (www.biorender.com). GraphPad Prism was employed for generating the plots. **Competing interest:** The authors have no conflicts of interest to declare. **Author contributions:** Conceptualization: T.S. and A.K.P. Methodology: A.K.P., S.C., R.K.N., and Y.K. Experimentation: T.S., S.T., R.P., S.K.G., V.B., V.K.N., M.P., P.D., and B.N.P Computational analysis: J.K. and S.C. Supervision: A.K.P. Writing-original draft: T.S. and A.K.P. Writing-review and editing: T.S., S.C., R.K.N., and A.K.P. The manuscript was read and approved by all the authors. **Material availability:** The corresponding author had full access to all the study data and made the final decision to submit it for publication. All necessary data to assess the conclusions in the paper are provided either in the paper itself or in the supplementary materials. The RNA-Seq datasets have been deposited in the Indian Nucleotide Data Archive (Accession ID: PRJEB66126).

