## Supplementary figures and table for "Phosphoglucomutase A mediated regulation of carbon flux is essential for antibiotic and disease persistence in *Mycobacterium tuberculosis*"

**Taruna Sharma[1,2] *et. al*.**

[1]Mycobacterial Pathogenesis Laboratory, Centre for Tuberculosis Research, Translational Health Science and Technology Institute, Faridabad, Haryana, India.

[2]Jawaharlal Nehru University, New Delhi, India.

[3]Experimental Animal Facility, Translational Health Science and Technology Institute, Haryana, India.

**#Correspondence-**

**Amit Kumar Pandey[1,3#] :**

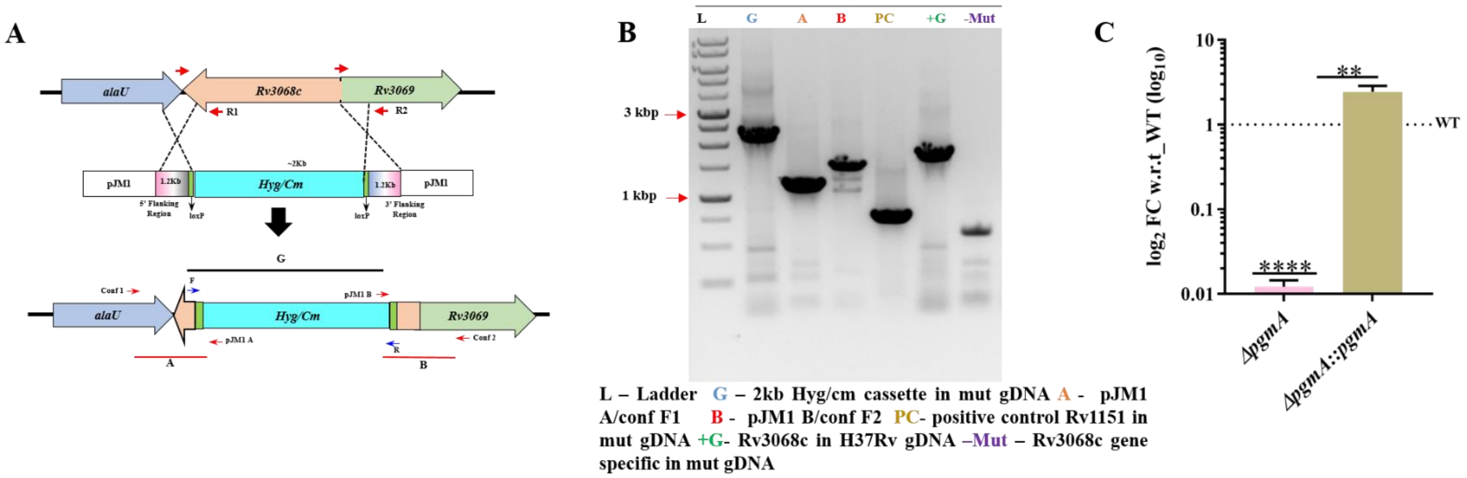


**Supplementary figure S1- Generation of Mtb *Rv3068c* deletion mutant** (A) *Rv3068c* (*pgmA*) mutant generation strategy by RecET mediated genetic recombineering *(1)* (B) *Rv3068c* mutant confirmation by PCR (C) Expression level of *pgmA* gene in *ΔpgmA* and *ΔpgmA::pgmA* by RT-qPCR and quantified by $\Delta\Delta C_T$ method. Statistical significance of (C) was determined using unpaired, non-parametric two-tailed t-test $^{**}P \leq 0.005$ and $^{****}P \leq 0.00005$. Data represent mean ± SEM (standard error mean) for technical triplicates.

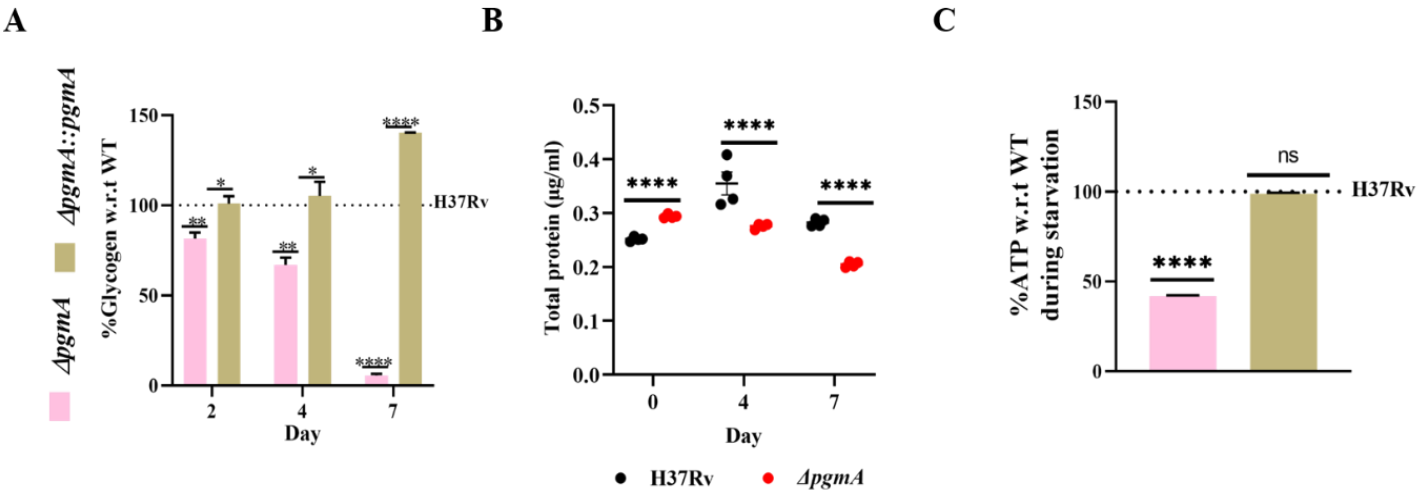


**Supplementary figure S2- Deletion of *pgmA* leads to impaired glycogen levels and overall biomass.** (A) Glycogen estimation in PBST using colorimetric based EnzyChrom glycogen assay kit (Bioassay Systems) (B) BCA estimation in PBST. (C) ATP measurement of H37Rv and *ΔpgmA* under starvation condition using BacTiter-Glo™ microbial cell viability assay kit (promega). Statistical significance of (A) to (C) was determined using unpaired, non-parametric two-tailed t-test *P ≤ 0.005, **P ≤ 0.005 and ****P ≤ 0.00005. Data represent mean ± SEM (standard error mean) for technical triplicates.

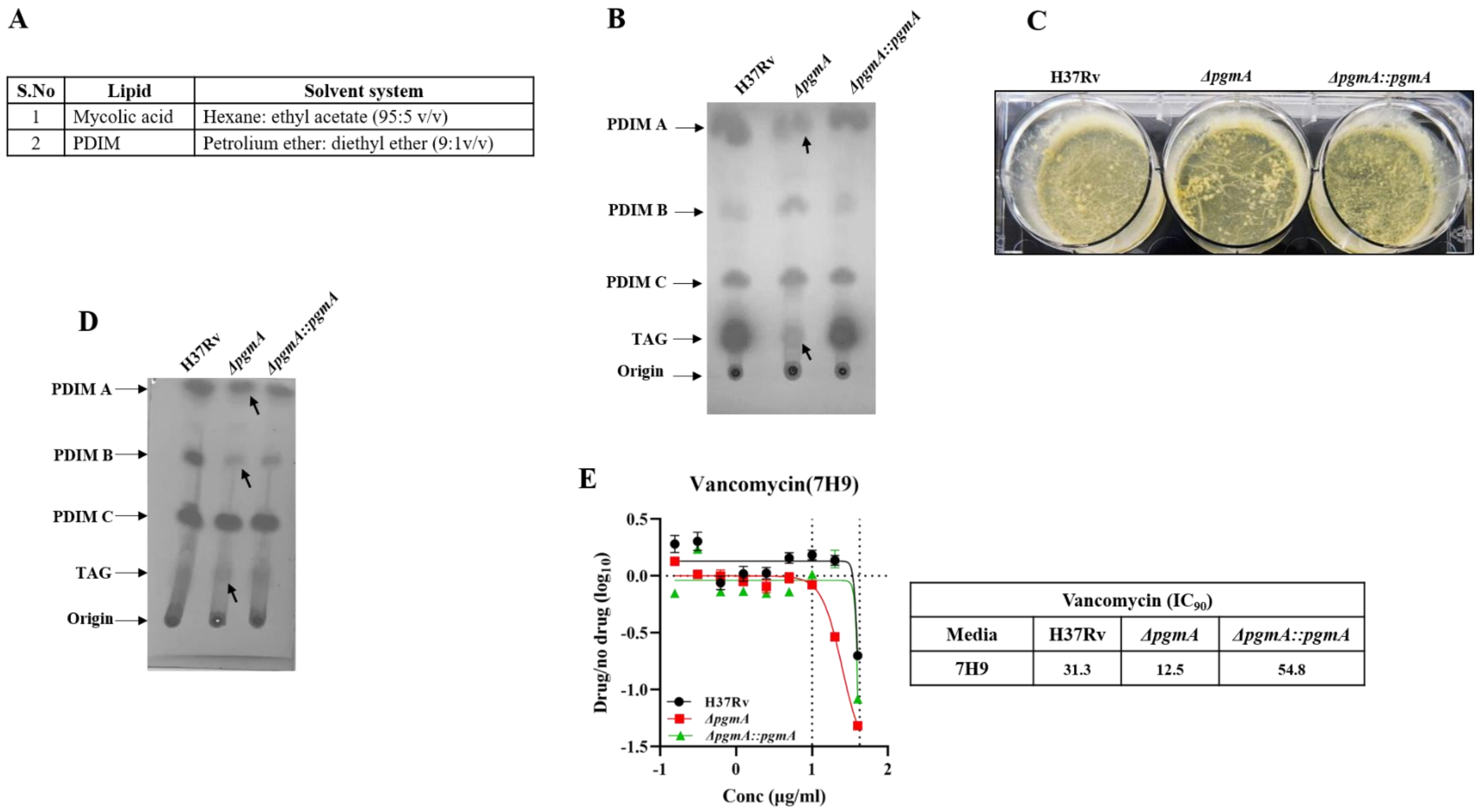


| S.No | Lipid | Solvent system |
|---|---|---|
| 1 | Mycolic acid | Hexane: ethyl acetate (95:5 v/v) |
| 2 | PDIM | Petrolium ether: diethyl ether (9:1v/v) |

| Vancomycin ($IC_{90}$) | | | |
|---|---|---|---|
| Media | H37Rv | *ΔpgmA* | *ΔpgmA::pgmA* |
| 7H9 | 31.3 | 12.5 | 54.8 |

**Supplementary figure S3- Deletion of *pgmA* impairs cellular integrity.** (A) Solvent system used for the separation of individual lipids by TLC from total lipids. (B) 1D separation of apolar fraction of total lipids by TLC. (C) *ΔpgmA* showing defect in biofilm formation. (D) TLC separation of apolar fraction of total biofilm lipids showing defects in PDIMs and TAGs. (E) MIC determination of vancomycin in H37Rv, *ΔpgmA* and *ΔpgmA::pgmA*. Abbreviations: PDIMs= phthiocerol dimycocerosates, TAGs=triacylglycerols, TLC= thin layer chromatography.

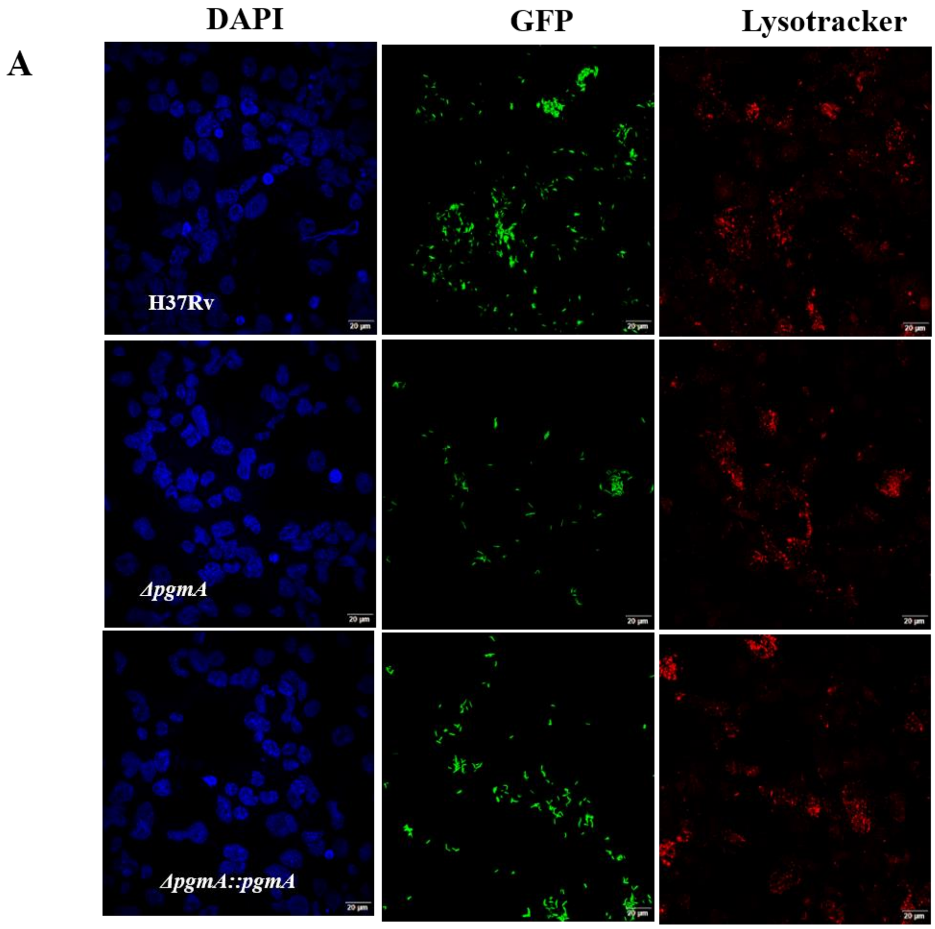


**Supplementary figure S4- THP-1 macrophages infected with Mtb** (A) Confocal imaging of THP-1 cells stained with DAPI (blue panel), bacteria having over-expressed GFP levels (Green panel) and lysotracker staining the acidic compartments (red panel). Images were captured using an Olympus FV3000 confocal microscope at 60× magnification (Scale bars, 20 μm).

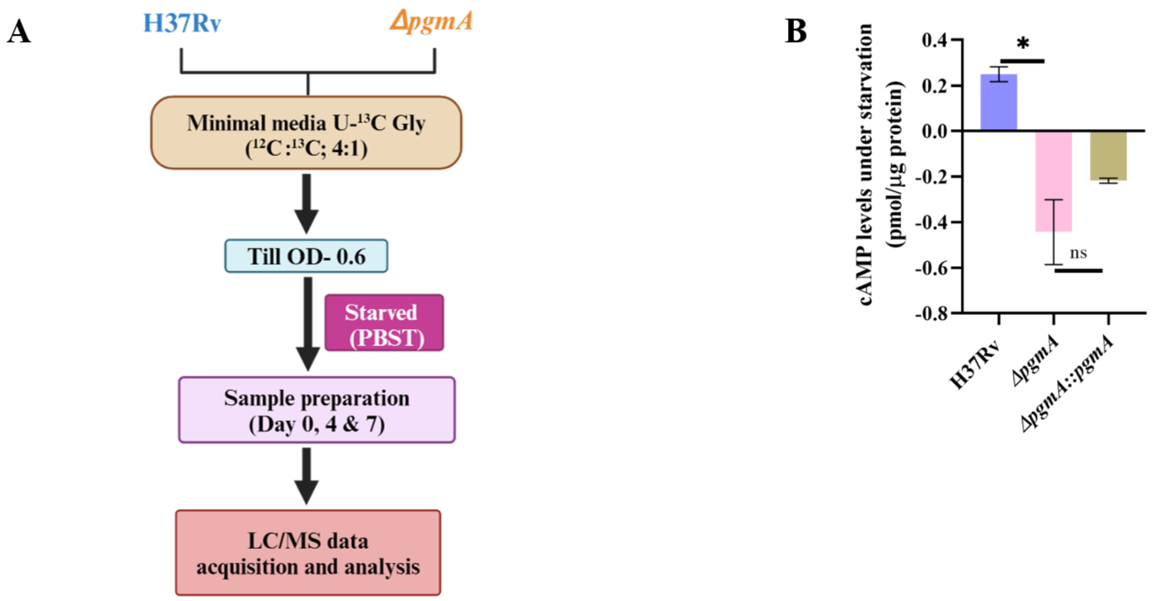


**Supplementary figure S5- *pgmA*-mediated regulation of carbon flux is crucial for the survival of Mtb under nutrient stress**. (A) Diagram illustrating the process of isotopic labeling with $^{13}C$, wherein Mtb strains were initially cultured in minimal media containing U-$^{13}C$ glycerol (0.1%, $^{13}C$ /$^{12}C$, ¼:v/v). Samples were collected on days 4 and 7 post-starvation and processed for liquid chromatography and mass spectrometry (LC/MS) data acquisition. (B) Under-representation of cAMP levels in *ΔpgmA* under starving conditions measured using direct immunoassay based kit. Statistical significance was determined using unpaired, non-parametric two-tailed t-test *P ≤ 0.005, ns=non-significant. Data represent mean ± SEM (standard error mean) for technical triplicates.

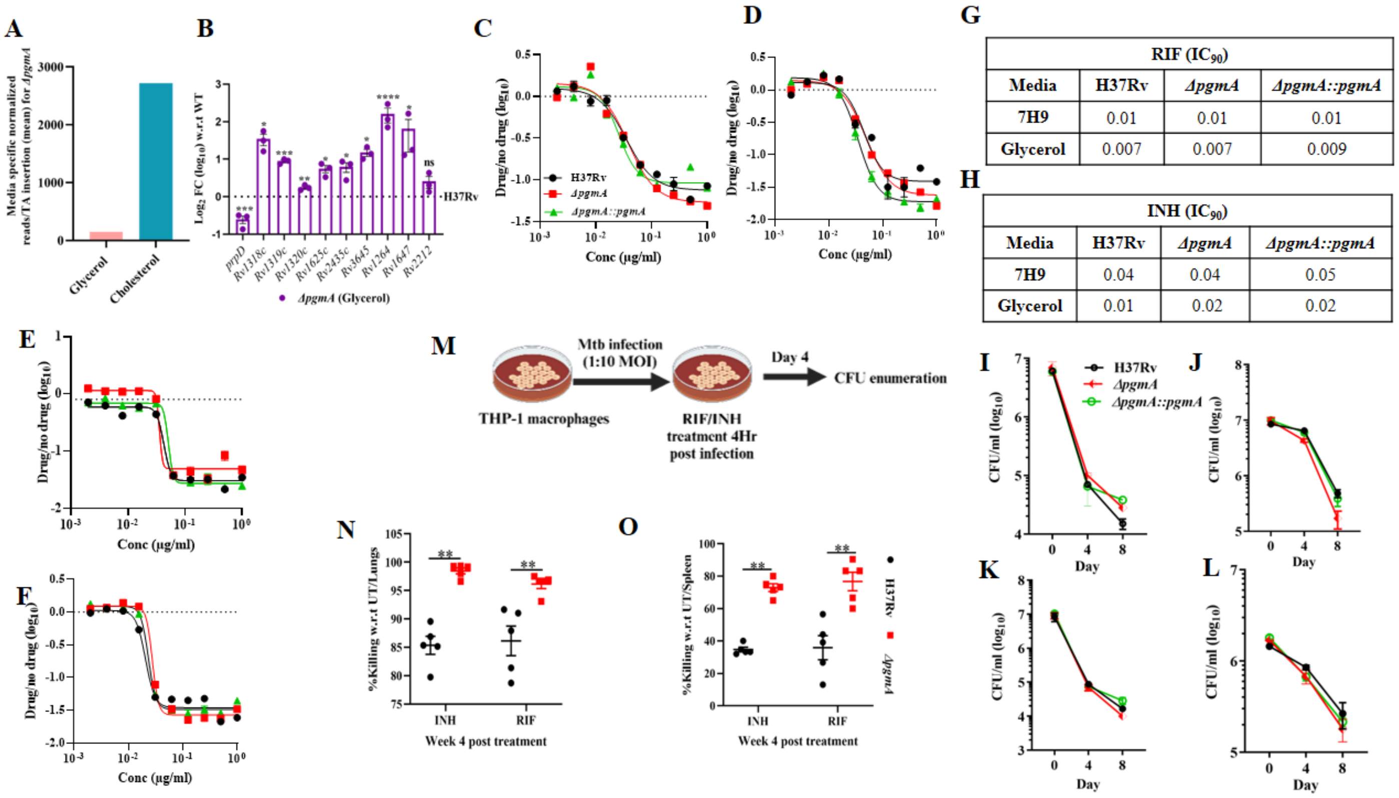


**G**

| RIF (IC90) | | | |
|---|---|---|---|
| Media | H37Rv | *ΔpgmA* | *ΔpgmA::pgmA* |
| 7H9 | 0.01 | 0.01 | 0.01 |
| Glycerol | 0.007 | 0.007 | 0.009 |

**H**

| INH (IC90) | | | |
|---|---|---|---|
| Media | H37Rv | *ΔpgmA* | *ΔpgmA::pgmA* |
| 7H9 | 0.04 | 0.04 | 0.05 |
| Glycerol | 0.01 | 0.02 | 0.02 |

**Supplementary figure S6- Data showing carbon source specific drug susceptibility of *ΔpgmA*.** (A)TraSH data depicting transposon insertion mutant of *pgmA* showing ~18-fold over-representation in cholesterol-rich media compared to glycerol-rich media. This calculation was based on the mean number of reads detected per TA insertion site (*2*). (B) The expression levels of adenylate cyclases under glycerol-specific media conditions in *ΔpgmA* with respect to (w.r.t) H37Rv. Bioluminescence-based assays utilized to determine the MICs of RIF and INH in 7H9 media (C and E) and glycerol media (D and F) respectively. IC90 values of (G) RIF and (H) INH under respective media conditions. In (I) to (L), time kill assay was conducted *in-vitro* using RIF and INH, with (I) and (K) representing 7H9 media, and (J) and (L) representing glycerol media, respectively. (M) Schematic depiction of the *ex-vivo* drug susceptibility assay conducted in the THP1 cell line is as follows: THP1 cells were infected at 1:10 MOI with subsequent drug treatment administered after 4 hours of infection. In (N) and (O), Percentage killing with respect to (w.r.t) untreated group was evaluated in lung (N) and spleen (O) of infected mice at week-4 post-treatment. Statistical significance in (B) and (N) to (O) was determined using unpaired, non-parametric, two-tailed t-test $*P \leq 0.005$, $**P \leq 0.005$, $***P \leq$

0.0005 and Mann-Whitney test **$P \leq 0.008$ respectively. Data represent mean ± SEM (standard error mean) for technical triplicates. Abbreviation: MOI= multiplicity of infection, MIC= minimum inhibitory concentration.

| Functional category | Up regulated genes | Down regulated genes | Total Up regulated genes | Total down regulated genes |
|---|---|---|---|---|
| **Starvation/late stationary phase/NRP** | *Rv 3290, Rv2891, Rv0251c, Rv3288c, Rv2699c, Rv0120c, Rv2632c, Rv2161c, Rv1288, Rv1834, Rv0458* | *Rv2665* | 11 | 1 |
| **Immune modulation** | *Rv1804c, Rv2353c, Rv1813c* | *Rv2307c, Rv1146 (mmpl13b)* | 3 | 2 |
| **Transcription factors** | - | *Rv3160c, Rv0792c, Rv2327, Rv2250c, Rv1499, Rv3124(moaR1), Rv0232, Rv0144, Rv0081, Rv0818* | 0 | 10 |
| **Transposon/insertion elements** | - | *Rv3186, Rv1313c, Rv0795, Rv1036c, Rv1585c, Rv0920c, Rv2651c, Rv3844, Rv2812, Rv1573, Rv1586c, Rv1580c* | 0 | 12 |
| **Membrane protein/permease/lectins/Cell wall/peptidoglycan** | *Rv2340c, Rv1004c, Rv1999c* | *Rv0666, Rv0488, Rv0219, Rv1382, Rv2254c, Rv0680c, Rv2307c, Rv0463, Rv0048c, Rv2293c, Rv0289 (espG3), Rv2723, Rv2686c (ABC transporter), Rv2325c, Rv2403c (lppR), Rv1986, Rv2643 (arsC), Rv3454, Rv0473, Rv3821, Rv1541c, Rv1217c (ABC transporter), Rv1146 (MmpL13b), Rv1038c (esxJ), Rv1793 (esxN), Rv1914c, Rv2620c, Rv0588 (yrbE2B), Rv0008c, Rv0584* | 3 | 30 |
| **Antibiotic resistance** | *Rv1930c, Rv3614c* | *Rv0095c, Rv0856, Rv0071* | 2 | 3 |
| **Intermediary metabolism/respiration/DNA synthesis & repair** | *Rv0356c, Rv0223c, Rv1393c, Rv1834 (lipZ)* | *Rv1571, Rv0791c, Rv0693(mftC), Rv0331, Rv2622, Rv0591 (mce2C), Rv3352c, Rv0166 (fadD5), Rv2381c (mbtD), Rv0812, Rv0694(mftD), Rv1939, Rv3703c, Rv1204c, Rv3701c, Rv2678c(hemE), Rv1089A (celA2a), Rv0322 (udgA), Rv1538c (ansA), Rv1203c, Rv3685c (cyp135), Rv3026c, Rv2435c (adenylyl cyclase), Rv0594 (mce2F), Rv2153c (MurG), Rv1888c, Rv3588c (canB), Rv0816c (thiX), Rv2423, Rv0762c, Rv3468c, Rv1853, Rv0601c, Rv1263 (amiB2), Rv0470c (pcaA), Rv1681 (moeX), Rv2934 (ppsD), Rv1570 (bioD), Rv1302 (rfe), Rv3556c (fadA6), Rv3436c (glmS), Rv0614 Rv0766c (cyP123), Rv3010c (pfkA), Rv0674, Rv0224c, Rv3322c, Rv1820 (ilvG), Rv3097c (lipY), Rv1319c (adenylyl cyclase), Rv2384 (mbtA), Rv0202c (mmpl11), Rv2949c, Rv0695 (mftE), Rv2840c, Rv0587 (yrbE2A), Rv2948c (fadD2), Rv2932 (ppsB), Rv0521, Rv0673 (echA4), Rv0726c, Rv2065 (cobH), Rv0469, Rv1847, Rv0539, Rv0189c, Rv3176c (mesT), Rv3312c, Rv0197, Rv2982c (gpdA2), Rv3473c (bpoA, Rv3581c (ispF), Rv3470c, Rv0370c, Rv2766c (FabG5), Rv3704c (gshA), Rv2505c (fadD35), Rv0096, Rv1546, Rv0968, Rv2737c, Rv3466, Rv1850 (ureC), Rv1641 (infC), Rv3202c, Rv0629c (recD), Rv1537 (dinX), Rv3201c, Rv2132, Rv2836c, Rv2885c, Rv2404c (lepA), Rv0269c, Rv2839c (infB), Rv3394c, Rv1003, Rv2755c, Rv3687c (rsfB), Rv2948c (fadD22)* | 4 | 97 |
| **TA system** | *Rv3357* | *Rv2758c, Rv0626, Rv0596c, Rv0623 (vapB30), Rv2862c (vapB23), Rv2827c, Rv2274c (mazF8), Rv1990* | 1 | 8 |
| **PE-PGRS** | *Rv3590* | *Rv0832, Rv1452c, Rv2107* | 1 | 3 |
| **Conserved hypothetical** | *Rv1670* | *Rv0863, Rv2489c, Rv3098c, Rv3647c, Rv3142c, Rv1684, Rv3163c, Rv2023c, Rv3439c, Rv0378, Rv2134c, Rv2722, Rv0610c, Rv2086, Rv1191, Rv2183c, Rv1682, Rv2566, Rv3771c, Rv2433c, Rv2076c, Rv2807, Rv2644c, Rv1060, Rv0036c, Rv1518, Rv0021c, Rv1116A* | 1 | 28 |

**Supplementary Table 1S:** Functional separation of up and down regulated DEGs obtained in RNAseq analysis exclusively in *ΔpgmA using* Go-ontology.

| Subsystems | Up-regulated_genes | Down-regulated_genes | Total_Up | Total_Down | Total |
|---|---|---|---|---|---|
| **Membrane Metabolism** | - | *Rv0166, Rv2381c, Rv2384, Rv2505c (fadD35), Rv2932, Rv2934, Rv2948c (fadD2), Rv2949c, Rv2293c, Rv1546,* | 0 | 10 | 10 |
| **Cofactor and Prosthetic Group Biosynthesis** | - | *Rv0202c, Rv1570, Rv2065, Rv2678c, Rv3322c* | 0 | 5 | 5 |
| **Fatty Acid Metabolism** | - | *Rv0166, Rv0673, Rv2505c, Rv2948c, Rv3556c* | 0 | 5 | 5 |
| **Translation/DNA synthesis & repair** | - | *Rv1850 (UreC), Rv1853 (ureD), Rv3202c (RecF), Rv2737c, Rv3466, Rv1641, Rv3202c, Rv0629c, Rv1537, Rv3201c, Rv2132, Rv2836c, Rv2885c, Rv2404c, Rv0269c, Rv2839c, Rv3394c, Rv1003, Rv2755c, Rv3687c* | 0 | 20 | 20 |
| **Valine, Leucine, and Isoleucine Metabolism** | - | *Rv0189c, Rv1820, Rv3470c, Rv3556c* | 0 | 4 | 4 |
| **Alanine, Aspartate, and Glutamate Metabolism** | - | *Rv1538c, Rv3704c* | 0 | 2 | 2 |
| **Glycolysis/Gluconeogenesis** | *Rv0458* | *Rv3010c (pfk)* | 1 | 1 | 2 |
| **Arabinogalactan biosynthesis** | | *Rv1302, Rv3468c, Rv3436c (glmS)* | 0 | 2 | 2 |
| **Beta oxidation of unsaturated fatty acids** | | *Rv0673, Rv3556c* | 0 | 2 | 2 |
| **Glycerophospholipid metabolism** | | *Rv2982c, Rv3176c* | 0 | 2 | 2 |
| **Mycobactin biosynthesis** | | *Rv2381c, Rv2384* | 0 | 2 | 2 |
| **Other Amino Acid Metabolism** | *Rv0223c* | *Rv1263* | 1 | 1 | 2 |
| **Pyruvate Metabolism** | | *Rv0694, Rv3556c* | 0 | 2 | 2 |
| **Cholesterol degradation** | | *Rv3556c* | 0 | 1 | 1 |
| **Degradation/Utilization/Assimilation** | | *Rv3176c* | 0 | 1 | 1 |
| **Ergothioneine biosynthesis** | | *Rv3703c* | 0 | 1 | 1 |
| **Folate Metabolism** | | *Rv0812* | 0 | 1 | 1 |
| **Glycerolipid metabolism** | *Rv0458* | - | 1 | 0 | 1 |
| **L-alpha-aminoadipic acid (L-AAA) biosynthesis** | *Rv3290c* | - | 1 | 0 | 1 |
| **Lipid metabolism** | | *Rv3097c (lipY), Rv2932 (ppsB), Rv2766c (FabG5)* | 0 | 3 | 3 |
| **Miscellaneous** | | *Rv3588c* | 0 | 1 | 1 |
| **Mycolic acid pathway** | *Rv1288* | *Rv0470c (pcaA)* | 1 | 1 | 2 |
| **Nucleotide Salvage Pathway** | - | *Rv0322* | 0 | 1 | 1 |
| **Peptidoglycan Metabolism** | - | *Rv2153c* | 0 | 1 | 1 |
| **Polyprenyl Metabolism** | - | *Rv3581c* | 0 | 1 | 1 |
| **Redox Metabolism** | - | *Rv0021c, Rv0694 (mftC), Rv0693 (mftD), Rv0695 (mftE)* | 0 | 4 | 4 |
| **Respiration** | - | *Rv0766 (Cyp123 P450),* | 0 | 1 | 1 |
| **Metabolic signaling** | - | *Rv2435c (AC), Rv1319c (AC)* | 0 | 2 | 2 |

**Supplementary Table S2:** Pathway analysis of up and down regulated metabolic genes obtained in RNAseq analysis.
